# Aberrant FICD-mediated AMPylation drives α-Synuclein pathology and overall protein dyshomeostasis in dopaminergic neurons in Parkinson’s disease

**DOI:** 10.64898/2026.03.30.715195

**Authors:** Aron Koller, Laura Hoffmann, Alexandra Bluhm, Alina Schweigert, Yanni Schneider, Marie Andert, Tobias Becker, Friederike Zunke, Thomas G. Beach, Geidy E. Serrano, Steffen Roßner, Jürgen Winkler, Pavel Kielkowski, Wei Xiang

**Author notes:** Correspondence : Pavel Kielkowski, Wei Xiang. These authors contributed equally to this work (Pavel Kielkowski, Wei Xiang).

## Abstract

**Background:** Filamentation induced by cAMP domain-containing protein (FICD) is an endoplasmic reticulum (ER)-resident adenylyltransferase that catalyzes protein AMPylation, a post-translational modification. Although FICD-mediated AMPylation has been linked to the fine-tuning of proteostasis and neuronal integrity, its role in neurodegenerative diseases characterized by protein dyshomeostasis remains unclear. Parkinson’s disease (PD) is defined by dopaminergic neurodegeneration and aggregation of α-synuclein (aSyn) as a consequence of impaired protein homeostasis. We therefore investigated whether dysregulated FICD-mediated AMPylation contributes to PD pathogenesis.

**Methods:** We combined analyses of human post-mortem PD brain tissue with complementary models, including midbrain dopaminergic neurons derived from human induced pluripotent stem cells (hiPSCs) of a PD patient carrying an *SNCA* gene duplication and its isogenic gene dosage-corrected control line, transgenic mouse models of synucleinopathy, and an aSyn-overexpressing H4 neuroglioma cell model. Genetic and pharmacological modulation of FICD activity was integrated with multi-proteomic approaches, including chemical proteomics-based AMPylation profiling, stable isotope labelling with amino acids in cell culture-based global protein turnover analysis, and whole-proteome profiling to identify AMPylation-associated molecular pathways.

**Results:** FICD was preferentially expressed in dopaminergic neurons and was upregulated in *SNCA* duplication PD patient-derived neurons, as well as in the basal ganglia of PD post-mortem brains and synucleinopathy mice. Despite this overall increase, the proportion of FICD-expressing dopaminergic neurons was reduced under PD conditions, suggesting selective vulnerability of dopaminergic neurons to FICD. Mechanistically, FICD selectively AMPylated lysosomal proteins, thereby linking AMPylation to the regulation of degradative pathways. Moreover, hyperactivation of FICD-induced AMPylation triggered ER stress, impaired lysosomal function, reduced protein turnover, and ultimately promoted aSyn aggregation and apoptotic cell death. Importantly, pharmacological inhibition of AMPylation reversed aSyn pathology and neurite degeneration in PD patient-derived neurons.

**Conclusions:** We identify the pathological relevance of FICD-mediated AMPylation in PD-related neurodegeneration and its contribution to aSyn aggregation through a bidirectional interplay with aSyn pathology. Our findings support FICD-mediated AMPylation as a defining molecular switch regulating intracellular protein homeostasis in PD and highlight the FICD-AMPylation pathway as a potential therapeutic target for restoring aSyn pathology and mitigating disease progression.

## Background

Parkinson’s disease (PD) is the most frequent neurodegenerative movement disorder, characterized by progressive loss of dopaminergic neurons in the substantia nigra of the midbrain and the formation of Lewy bodies (LBs), primarily composed of aggregated α-synuclein (aSyn) (1). Via the assembly of monomers into less soluble aggregates, aSyn acquires toxic properties that drive neurodegeneration. Although the causal trigger leading to aSyn aggregation remains unclear, disturbed protein homeostasis is increasingly recognized as a key pathogenic mechanism (2).

Protein homeostasis depends on precisely tuned pathways controlling protein synthesis, folding, trafficking, and degradation. In the endoplasmic reticulum (ER), the unfolded protein response (UPR) orchestrates protein folding, while the autophagy-lysosome pathway and the ubiquitin-proteasome system mediate clearance (2). For maintaining cellular protein equilibrium, protein post-translational modifications (PTMs) play a crucial role through fine-tuning the stability, localization, and activity of modified proteins. Among PTMs, AMPylation has recently emerged as an important regulator of protein homeostasis (3–5) and neurodevelopment (6).

AMPylation involves the covalent addition of adenosine 5′-*O*-monophosphate (AMP) to serine, threonine or tyrosine residues of substrate proteins. AMPylation-catalyzing enzymes, adenylyltransferases, were initially described in bacteria, where Fic (filamentation induced by cAMP) domain-containing proteins were shown to AMPylate host proteins during infection (7, 8). In humans, FICD is currently the sole known Fic-domain-containing enzyme. The ER-resident FICD protein functions as both an AMP-transferase and de-AMPylase. Its best-characterized substrate is the ER chaperone binding immunoglobulin protein (BiP, also known as HSPA5), a central regulator of the UPR, whose AMPylation state dynamically calibrates ER protein folding capacity and downstream apoptotic signaling (9).

Recently, genetic FICD variants or mutations have been associated with neurological disorders, such as hereditary spastic paraplegia (10) and infancy-onset diabetes with severe neurodevelopmental delay (11). Moreover, emerging evidence implicates the FICD-AMPylation pathway in PD-related mechanisms. In *Caenorhabditis elegans*, AMPylation modulates aSyn aggregation and toxicity (12). In rodents, FICD is highly expressed in the substantia nigra, the brain region severely affected in PD (13).

Studies of AMPylation are technically challenging due to its low abundance. Yet recent advances in chemical proteomics now enable profiling AMPylation targets (14–17). We recently established a chemical-proteomic approach for high-throughput AMPylation detection in living cells (6), using a metabolically activatable adenosine analog (pro-N6pA) that allows intracellular labeling, enrichment, and mass spectrometric identification of AMPylated proteins. This strategy revealed multiple AMPylated proteins, including cysteine cathepsins, SQSTM1, and PLD3, key proteostatic regulators that are closely linked to neurodegenerative diseases (18–20).

Given the emerging evidence implicating the FICD-AMPylation pathway in maintaining protein homeostasis and neuronal integrity, we hypothesize that this pathway contributes to neurodegenerative processes in PD. Using rodent models, human *post-mortem* brains, as well as midbrain neurons derived from human induced pluripotent stem cells (hiPSCs), we show that FICD is preferentially expressed in dopaminergic neurons. We also demonstrate consistently upregulated FICD expression in the basal ganglia of PD *post-mortem* brains and corresponding transgenic mouse models, as well as in PD patient-derived dopaminergic neurons. By modulating both aSyn expression and FICD activity in combination with multiple proteomic approaches, we demonstrate that FICD selectively AMPylates lysosomal proteins. Hyperactivation of FICD-mediated AMPylation disrupts the ER-lysosome axis of protein homeostasis, increases aSyn aggregation, and promotes apoptosis. Importantly, pharmacological inhibition of FICD-mediated AMPylation mitigates aSyn aggregation and restores neurite integrity in PD patient neurons.

## Methods

### Cell culture

#### H4 cells

H4 human neuroglioma cells (ATCC, HTB-148) stably expressing human wildtype (WT) aSyn (H4-aSyn) and control cells (H4-Ctrl) were generated using lentiviral vectors pCMV::aSyn-RES-GFP and pCMV::RES-GFP, respectively, as described previously (21). H4 cells were cultured in Opti-MEM™ Reduced Serum Medium GlutaMAX™ Supplement (Thermo Fisher Scientific) supplemented with 10% FBS (Sigma Aldrich) and 1% penicillin-streptomycin (Thermo Fisher Scientific).

#### Transfection of H4 cells

Twenty-four hours prior to transfection, 2 × 10^5^ or 4 × 10^4^ H4 cells were seeded per well in a 6-well or 24-well-plate, respectively. Transfections were performed using Fugene4K (Promega) and expression vectors encoding human FICD variants, including wildtype (WT) FICD (FICD-WT), the inactive mutant FICD-H363A, and the constitutively active AMPylase FICD-E234G at a 1:1 (v/w) ratio according to manufacturer’s instructions. For transfections in 6- and 24-well-plates, 2 µg and 0.5 µg DNA was used respectively. Mock transfections were performed with the corresponding empty expression vector at equal DNA amounts.

#### Human iPSCs

The hiPSC SNCA^Dupl^ line (UKERi7GG-S1-004) was generated from dermal fibroblasts obtained via the skin biopsy from a PD carrying heterogenous *SNCA* duplication. (22, 23) The biopsy was collected at the Movement Disorder Clinic, Department of Molecular Neurology, University Hospital Erlangen. All procedures involving the generation and use of hiPSCs were approved by the local Institutional Review Board (Nr. 259_17B). The isogenic hiPSC SNCA^Ctrl^ line (UKERi7GG-S1-004-1) were generated by CRISPR-Cas9 gene editing of the *SNCA* duplication hiPSC line SNCA^Dupl^ (UKERi7GG-S1-004) as described previously (22).

#### Differentiation of hiPSC-derived midbrain neurons

Midbrain neurons were generated from hiPSCs via an embryoid body stage and neural precursor cells (NPCs) using a small-molecule based protocol (24), as modified in our previous work (22, 23). Briefly, NPCs were cultured in NPC medium, consisting of N2B27 (50% Dulbecco’s modified Eagle’s medium/F12 GlutaMAX, 50% Neurobasal medium, 1:200 N2, 1:100 B27, 1:100 penicillin-streptomycin) supplemented with 3 µM Chir, 0.5 µM purmorphamine (PurMA), 10 µM SB531542, 150 µM ascorbic acid. Cells were split in ratios of 1:3 - 1:5 onto new 12-well plates for NPC maturation. To induce midbrain differentiation, NPC medium was replaced by N2B27 medium containing 100 ng/mL FGF-8, 1 µM PurMA, and 200 µM ascorbic acid (differentiation medium). Cells were cultivated in differentiation medium until the 8th day of differentiation and medium was re-newed every other day. On day 8, differentiation medium was replaced by medium containing 10 ng/mL BDNF, 10 ng/mL GDNF, 1 ng/mL TGF-β3, 500 µM cAMP, 0.5 µM PurMA, and 200 µM ascorbic acid (maturation medium). On day 9, 1 × 10^6^ cells were plated in 12-well plates or 3 × 10^5^ cells were seeded onto coverslips in 24-well plates in maturation medium supplemented with 10 µM ROCK inhibitor and removed after additional 24h. On day 11, PurMA was withdrawn from maturation medium, and cells were cultivated in this medium until the day 24, replacing maturation medium every other day.

### Human *post-mortem* tissue

#### Substantia nigra brain tissue

Case recruitment and autopsy were performed in accordance with guidelines effective at the Arizona Study of Aging and Neurodegenerative Disorders and Brain and Body Donation Program (25). The required consent was obtained for all cases. Clinical data of the cases are shown in Table S1. Transverse midbrain sections (40 µm) comprising the substantia nigra at the level of the red nucleus, exit of the oculomotor nerve and superior colliculus were used for immunohistochemistry.

#### Putamen brain tissue

*Post-mortem* putamen tissue was obtained from the Netherlands Brain Bank (NBB, Netherlands Institute for Neuroscience, Amsterdam, http://www.brainbank.nl). PD diagnoses were established clinically based on characteristic motor symptoms (bradykinesia, rigidity, tremor) and confirmed neuropathologically by the presence of Lewy body (LB) pathology. All PD cases included in this study exhibited neocortical LB pathology corresponding to Braak Lewy stages 5 - 6 (Table S2). Putamen tissue was used for Western blot analysis.

### Rodent brains

Transgenic mice overexpressing human aSyn under the Thy1 promoter (Thy1-aSyn, Line 61) or the MBP promoter (MBP-aSyn, Line 29) were generated as described previously (26, 27). Both animal models were assessed at a symptomatic stage (Thy1-aSyn, male, 12-month-old; MBP29-aSyn, female, 2-month-old) and compared to their sex, and aged-matched non-transgenic litter mates. For FICD distribution analysis, Sprague-Dawley WT rats aged 10 - 12 months of mixed sex were used. Breeding and housing of these animals as well as brain preparation were approved by the local governmental administrations for animal health (TS03-20, Division for Animal Welfare, Friedrich-Alexander-University Erlangen-Nürnberg, and AZ. 55.2,2-2532-2-1489, AZ. 55.2-DMS 2532-2-218 Regierung von Unterfranken, Würzburg).

### Western blot

Cells were lysed using RIPA buffer (50 mM Tris/HCl, pH 8.0, 150 mM NaCl, 5 mM EDTA, 1% NP40, 0.5% sodium deoxycholate, 0.1% SDS) containing 1× protease inhibitor cocktail (PIC) (Roche) for 30 min on ice. Brain tissue was first homogenized in a buffer containing 50 mM Tris/HCl pH 8.0, 150 mM NaCl) using a Braun Potter S Homogenizer (Sartorius AG) and then mixed with 4× RIPA in a 1:3 ratio, followed by incubation on ice for 30 min. Next, samples were centrifuged at 10,000 g for 10 min at 4°C and protein content of the supernatant was determined using the Pierce™ BCA Protein Assay Kit (Thermo Fisher Scientific). Samples were brought to the same concentration and boiled in LDS-sample buffer with reducing agent (Thermo Fisher Scientific) at 70°C for 10 min or in 5× Laemmli buffer containing 0.3 M Tris/HCl pH 6.8, 50% glycerol, 1% SDS, 0.05% bromophenol blue, and 5% 2-Mercaptoethanol at 99°C for 10 min. For SDS-PAGE, 20-40 µg protein was loaded on either NUPAGE 4 - 12% Bis-Tris gels (Thermo Fisher Scientific) with MES buffer or selfcast 12% acrylamide gels with Tris/glycine buffer and separated by electrophoresis run at 120 V for 90 min. For protein transfer, the gel was blotted on a PVDF Fl membrane (Merck Millipore) and fixed in 4% paraformaldehyde (PFA) for 15 min. The membranes were blocked in Intercept Blocking Buffer (Li-Cor) for 1 h and incubated overnight with primary antibodies (Table S3) at 4°C. After washing three times with TBS containing 0.1% Tween20, membranes were incubated with fluorophore-coupled, species-matched secondary antibodies (Table S3) for 1 h at room temperature (RT). Immunosignals were detected using a Fusion FX7 imaging system (PeqLab) or an Odyssey M imaging system (Li-Cor). Quantification of the relative levels of target proteins was performed using Image StudioLite Software (Version 5.2.5, Li-Cor) or Empiria Studio (Version 3.3, Li-Cor). For each experiment, the signal intensity of proteins of interest was normalized to the corresponding signal of the designated loading control, such as immunointensity of glyceraldehyde-3-phosphate-dehydrogenase (GAPDH), total protein stain using Revert 520 Total Protein Stain (Li-Cor), or Coomassie Brilliant Blue G 250 (Bio-Rad) on the SDS-PAGE gel.

### CTSB and CTSD activity assay

The measurement of the activity of cathepsins, CTSB and CTSD, from cell lysates was performed as previously described (19). In short, cells were lysed in acidic buffer (50 mM sodium acetate, 0.1 M NaCl, 1mM EDTA, 0.2% Triton X-100, pH 4.5) and centrifuged at 13,000 g for 15 min at 4°C. 2 µL supernatant was incubated in 98 µL acidic buffer containing 20 µM and 10 µM quenched fluorogenic peptides (BML-P137 and BML-P145, Enzo) for CTSB and CTSD, respectively, for 30 min at 37 °C. Enzymatic activity was measured using a ClarioStar (BMG LABTECH) plate reader (ex: 365 nm, em: 440 nm for CTSB and ex: 322, em: 381 for CTSD). All values were corrected for background fluorescence and normalized to protein content.

### Reverse transcription quantitative polymerase chain reaction (RT-qPCR)

Cells were pelleted in phosphate buffered saline (PBS) at 300× g for 5 min at 4°C. Total RNA was extracted using the RNeasy Kit (Qiagen). RNA concentration was measured with a NanoDrop spectrophotometer and 400 ng of RNA was reverse transcribed into cDNA using the DNA GoScript™ Reverse Transcription System (Promega). qPCR was performed on a LightCycler 480 (Roche) using the SsoFast™ EvaGreen® Supermix (Biorad) with primers (Table S4). Relative levels of target mRNA were determined with the LightCycler480 Software (Roche) and normalized to 18S and GAPDH mRNA.

### aSyn aggregation assays

#### Solubility assay

Cells collected from two wells of a 6-well plate were harvested and homogenized in 50 mM Tris/HCl pH7.4, 175 mM NaCl, 5 mM EDTA, 1% Triton X-100 using a Braun Potter S Homogenizer (Sartorius AG). After 30 min incubation on ice, protein concentration of the homogenate was determined, adjusted to 1 µg/µl and 70-100 µg of protein was ultracentrifuged at 100,000 g for 60 min at 4°C. The pellet was resuspended in half of the volume of the supernatant with a buffer containing 0.5 M Urea and 5% SDS and 5× Laemmli buffer was added to both fractions, followed by incubation at 98°C for 10 min. Lastly, samples were submitted to electrophoresis using Tris/glycine buffer with selfcast 12% acrylamide gels at 120V for 90 min and blotted on a PVDF-Fl membrane.

#### Filter trap assay

Cells from two wells of a 6-well plate were harvested, resuspended in 100 µL PBS containing PIC and sonicated three times and centrifuged at 1000 g for 5 min at 4°C. After determining the protein concentration of the supernatant, 30 µg protein was mixed with 80 µL 0.1% SDS in H_2_O and PBS for a total volume of 120 µL and filtered through a 0.2 µm cellulose acetate membrane (Sterlitech) for 15 min using a vacuum supported dot bot chamber (Minifold SRC-96/1, Schleicher & Schuell). Before and after filtration, the membrane was washed twice with PBS. To control for equal protein loading, a nitrocellulose membrane (0.2 µm, Cytiva) was placed beneath the cellulose acetate membrane and stained for total protein with Ponceau S solution (Sigma Aldrich). The cellulose acetate membrane was fixed with 4% PFA for 10 min, washed with PBS (3×) and blocked with Intercept Blocking Buffer (Li-Cor) for 1 h before proceeding with immunodetection as described in the “Western blot” section.

### Flow cytometry analysis

Apoptosis was analyzed using an Annexin V / propidiumioide (PI) assay (Biolegend) according to the manufacturer’s instructions. As a positive control and for gating setup, cells were treated with 1 µM staurosporine (Cell Signaling Technology) for 2 h prior to harvesting. For the Annexin/ PI assay, 2 × 10^5^ cells were harvested with Accutase (Sigma) and pelleted at 300 g for 5 min. After washing with PBS, cells were resuspended in 150 µL Annexin V binding buffer (BB) and 50 µL Annexin V-APC (1:100 in BB). After incubation for 7 min, 10 µL PI (1:200 in BB) was added and after 3 min of incubation time, the cells were diluted in 200µl BB and analyzed with the BD LSRFortessa™ Cell Analyzer (BD Biosciences GmbH). Quantification was performed using FlowJo™ Software (BD Biosciences GmbH).

### Immunocytochemistry

#### Fluorescent immunostaining

Cells cultured on coverslips were treated with 2% PFA for 10 min, followed by 4% PFA for 10 min. After three PBS washes, cells were blocked for 60 min with fish skin gelatin buffer (TBS with 0.4% cold water fish gelatin in water, 1% BSA and 0.1% Triton X-100). Subsequently, cells were incubated with a primary antibody (Table S3) overnight at 4°C followed by incubation with a species-matched, fluorescence-labelled secondary antibody (Table S3) for 1 h at RT. Finally, cell nuclei were counterstained with DAPI (Sigma, 1:10,000 in PBS) for 7 min at RT and mounted using ProLong Gold solution (Thermo Fisher Scientific). Images were captured using an Axio Observer inverted fluorescence microscopes (Carl Zeiss) or a confocal laser scanning microscope (LSM 780, Carl Zeiss).

#### aSyn subcellular distribution

The distribution of aSyn within neuronal somata and neurites was evaluated using Zen 2013 software (Carl Zeiss). For each neuron positive for βIII-tubulin (TUBB3), 1 - 3 neurites were randomly selected, and cytosolic as well as neuritic regions were delineated to quantify aSyn signal intensity in both compartments.

#### FICD expression in hiPSC-derived dopaminergic neurons

For analysis of the proportion of FICD expressing and non-expressing neurons within dopaminergic (TH+/TUBB3+) and non-dopaminergic (TH-/TUBB3+) neurons were counted across five randomly selected regions per coverslip using Zen 2013. For determination of FICD intensity, 5 - 9 TUBB3+ neurons expressing either FICD alone or both FICD and TH were analyzed from five region per coverslip by delineating cytoplasmic region of the neuronal soma.

#### Neurite morphology

Neurite diameter and volume were measured with IMARIS (Bitplane) using the software’s filament tracing function to automatically reconstruct neurites within a randomly defined image region. Diameter analysis was restricted to primary neurites prior to the first branching point, whereas volume measurements included neurite segments from the first through the fifth branching point.

### Immunohistochemistry

#### Immunofluorescence staining in rat brain sections

Free-floating brain sections were washed (3× 5 min in TBS) and antigen retrieval was performed in citrate buffer (1.8 mM citric acid and 8.2 mM sodium acetate) for 30 min at 80°C, followed by 1 h blocking using TBS containing 3% donkey serum and 0.3% Triton X-100. Sections were stained with primary antibodies (Table S3) overnight at 4°C, washed and incubated with fluorophore-coupled, species-matched secondary antibodies (Table S3), for 1 h at RT. After three TBS washes, sections were counterstained with DAPI (1:10,000 in TBS) for 10 min, washed, and mounted on glass slides using ProLong Gold mounting solution.

#### Single labelling of FICD in human brain sections

Single labelling immunohistochemistry (DAB-Ni) was performed on free-floating transverse sections of the midbrain (40 µm) of human *post-mortem* tissue. Sections were washed in PBS and endogenous peroxidases were inactivated with 60% methanol containing 1% H_2_O_2_ for 1 h. Unspecific binding was then blocked in PBS-T containing 2% BSA, 0.3% milk powder and 0.5% normal donkey serum before incubating brain sections in the same solution with the primary anti-FICD antibody for 40 h at 4°C. Subsequently, sections were washed in PBS-T (3× 7 min) and then incubated with secondary biotinylated donkey anti-rabbit antibody (Dianova; 1:1,000) in a mixture of blocking solution and PBS-T (1:2) for 1 h at RT. After three washes, the sections were incubated with ExtrAvidin-peroxidase (Sigma; 1:1,500 in PBS-T) for 1 h. Binding of peroxidase was visualized by incubation with 1 mg 3,3’-Diaminobenzidin (DAB), 20 mg nickel ammonium sulfate and 2.5 µL 30% H_2_O_2_ per 5 mL 0.5 M Tris/HCL buffer (pH 8.0) for 3 min. Finally, sections were mounted onto glass slides and cover slipped. Immunohistochemically stained sections were examined with an Axio-Scan.Z1 slide scanner connected with a Colibri.7 light source and a Hitachi HV-F202SCL camera (Carl Zeiss). Images were digitized by means of ZEN 2.6 software and analyzed using the ZEN 3.6 imaging tool.

For each individual case, the area of the substantia nigra pars compacta was delineated according to the distribution of the nigral matrix and nigrosomes, respectively, enclosing pigmented NM-containing neurons. Analysis of FICD presence was performed using trainable segmentation with Apeer Machine Learning toolkit and ZEN blue 3.8 Intellesis software (Zeiss).

### Treatment with closantel

For the treatment of H4 cells, 10 µM closantel (Sigma-Aldrich) was added to the media at the start of transfection with FICD plasmids. For hiPSC-derived midbrain neurons, 10 µM closantel was added to the maturation medium at day 23 of the neuronal differentiation. Cells were either treated with closantel or the corresponding DMSO control for 30 h, after which the medium was removed and cells were washed with PBS.

### Mass spectrometry-based proteomics

#### Liquid chromatography-tandem mass spectrometry (LC-MS/MS) measurement

LC-MS/MS analyses were performed on an Orbitrap Eclipse Tribrid Mass Spectrometer, interfaced with an UltiMate 3000 Nano-HPLC system via a Nanospray Flex ion source and a FAIMS Pro interface (all from Thermo Fisher Scientific). Peptide samples were first loaded onto an Acclaim PepMap 100 μ-precolumn cartridge (5 μm, 100 Å; 300 μM ID × 5 mm, Thermo Fisher Scientific), followed by chromatographic separation at 40°C using a PicoTip emitter (non-coated, 15 cm length, 75 μm ID, 8 μm tip; New Objective) packed in-house with Reprosil-Pur 120 C18-AQ material (1.9 μm particle size, 150 Å pore size; Dr. A. Maisch GmbH). LC buffers comprised MS-grade water (A) and acetonitrile (B), both supplemented with 0.1% formic acid. The short gradient ran from 4-35.2% B over 60 min (0-5 min: 4%, 5-6 min: 7%, 7-36 min: 24.8%, 37-41 min: 35.2%, 42-46 min: 80%, 47-60 min: 4%) at a flow rate of 300 nL/min.

#### Data-dependent acquisition (DDA)

DDA was used for analysis of AMPylation enrichment in H4 cells as described in our previous work (28).

#### Data-independent acquisition (DIA)

DIA was used for the analysis of the whole proteome of H4 cells and AMPylation enrichment from NPCs and midbrain neurons. Field asymmetric ion mobility spectrometry (FAIMS) was performed using a single compensation voltage (CV) of -45V. The mass spectrometer operated in DIA mode, with each cycle comprising one MS1 scan followed by 30 MS^2^ scans. MS^1^ spectra were acquired in positive ion mode with an Orbitrap resolution of 60,000, using a standard AGC target and a maximum injection time of 50 ms. The scan range was typically set to m/z 200-1800, or m/z 120-1700 where indicated, and the RF Lens was operated at 30%. The precursor mass range was defined as m/z 500-740, with an isolation window of 4 m/z and an overlap of 2 m/z between windows. MS^2^ spectra were recorded with an Orbitrap resolution of 30,000, an AGC target of 200%, and an automatic maximum injection time. Higher-energy collisional dissociation (HCD) was applied at a normalized collision energy of 35%, and the RF Lens remained at 30%. The MS^2^ scan range was set to automatic.

#### Volcano plots generation and statistics

Raw intensities were imported to Perseus 2.0.9.0 (29). Quantified values were log2-transformed and samples were annotated and proteins with fewer than three valid values per group were removed. Next, missing values were imputed from a normal distribution. For differential analysis, a two-sided two-sample Student’s t test was performed with Benjamini-Hochberg adjustment across all proteins (FDR = 0.05). For visualization, log2(fold change) (probe/control) and -log10(p-value) were exported to Python 3.11.7 and volcano plots were generated with matplotlib (30).

#### Hierarchal clustering analysis

Hierarchal clustering was performed in Perseus (version 2.0.11.0). For the SILAC experiment, protein half-lives from the 938 commonly detected proteins with corresponding half-lives were used for the analysis. For the whole proteome data, one-way ANOVA was performed (FDR = 0.05, 250 randomizations, S_0_ = 0) and significant proteins were used in downstream analyses. Significant proteins were Z-scored and subjected to unsupervised hierarchical clustering using Euclidean distance and average linkage (300 clusters, 10 iterations, 1 restart). For subclustering analysis of the whole proteome data, results from hierarchal clustering were exported to Python 3.12 and analyzed using SciPy (31), seaborn (32), and matplotlib (30) packages.

#### Pathway overrepresentation analysis

For the chemical proteomics analysis, gene hits (p < 0.01, -log2(pro-N6pA/ DMSO) > 1) were submitted to ShinyGO v0.85 (organism: Homo sapiens) (33). For the whole proteome data, clustered proteins from hierarchal clustering analysis were used. The KEGG pathway library was tested against the organism-specific default background. Enrichment p-values were computed by hypergeometric testing and adjusted for multiple comparisons using Benjamini-Hochberg FDR control. Pathways with FDR < 0.05 were considered significant. Results were filtered by FDR and ranked by enrichment fold change.

#### Whole proteome analysis using SP3 workflow

The whole proteome analysis and preparation of SILAC sample was done using the SP3 protocol (34) utilizing a liquid handling robot (Hamilton Microlab Prep) as described in our previous work (22).

#### Chemical proteomics – enrichment of AMPylated proteins using SP2E workflow

The phosphoramidate probe pro-N6pA was synthesized as described previously (6). Probe treatment was performed by applying 100 µM pro-N6pA for 16 h at 37°C. DMSO was used as the vehicle control. For NPCs, the probe was added to fresh NPC medium, when cells reached 80% confluency. For neurons, the probe was added to N2B27 medium on day 23 of the midbrain differentiation. After 16 h, medium was removed and cells were washed with PBS. The suspension was centrifuged at 300 g for 5 min, and cell pellets were snap-frozen and stored at -80°C until further processing. Next, cells were lysed using 80-100 μL of lysis buffer (1% NP-40, 0.2% SDS, 20 mM Hepes, pH 7.5) via rod sonication (10s, 20% intensity). Lysates were centrifuged at 13,000 g at 4°C for 10 min and the protein concentration of the supernatant was performed with a Pierce™ BCA Protein Assay Kit.

Enrichment of the AMPylated proteins from H4-cells was performed in the large-scale format starting from 400 μg protein as described previously (28). Enrichment of modified proteins from NPCs (100 µg) and midbrain neurons (30-40 µg) using the small-scale SP2E workflow was conducted in a 96-well plate format. Initially, in-cellulo probe-labeled lysates (or DMSO control) were diluted to a total volume of 40 μL using lysis buffer. The enrichment was performed by a liquid handling robot (Hamilton Microlab Prep) as described previously (28). After the tryptic digest, the beads were washed manually with 20 μL of TEAB buffer (50 mM in mass spectrometry (MS)-grade water) and 20 μL of 0.5% MS-grade formic acid (in MS-grade water), each time incubating at 40°C and 600 rpm for 5 min. The combined eluates were supplemented with 0.9 μL MS-grade formic acid, placed on a magnetic stand for 15 min, and transferred to fresh 1.5 mL tubes. Following an additional 2 min magnetic separation, the final supernatants were transferred to MS vials to ensure removal of any residual beads and then subjected to LC-MS/MS analysis.

### SILAC (stable isotope labeling with amino acids in cell culture) proteomics

#### Pulse-chase experiment

Dulbecco’s modified Eagle’s medium (DMEM)-SILAC (Thermo Scientific) lacking arginine and lysine was supplemented with 1% penicillin-streptomycin and stable isotope-labelled arginine and lysine. The “light” SILAC medium was supplemented with L-[^12^C_6_,^14^N_2_] lysine and L-[^12^C_6_,^14^N_4_] arginine (Sigma Aldrich), whereas the “heavy” SILAC medium contained L-[^13^C_6_,^15^N_2_] lysine and L-[^13^C_6_,^15^N_4_] arginine (Sigma Aldrich). Both media were supplemented with L-[^12^C_5_,^14^N] proline and 10% dialyzed FBS. Prior to the pulse-chase experiment, H4 cells were cultivated for 14 days in light or heavy SILAC medium, after which full incorporation of the respective isotopes was confirmed via LC-MS/MS. For the pulse-chase experiment, H4-cells were transiently transfected with FICD variants for 24 h until the label switch was performed. Specifically, cells that were previously cultivated in light SILAC medium, were “pulsed” with heavy SILAC medium and vice versa. For 2, 8, and 24 h the cells were “chased” and harvested at the respective time points. The cell pellets were snap-frozen in liquid nitrogen and stored at -80°C until further processing.

#### SILAC data analysis

SILAC-based proteomic data were processed using MaxQuant (version 2.6.5.0.) with the search type set to MaxDIA. Multiplicity was configured for two labels corresponding to heavy isotopes of arginine (Arg10) and lysine (Lys8). Trypsin was selected as the proteolytic enzyme, and label-free quantification (LFQ) was enabled without using an internal isotope standard. The main output file used for downstream analysis was proteinGroups.txt.

The resulting MaxQuant output was imported into Perseus (version 2.0.11.0) for further analysis. Beforehand, zero values were replaced with NaN to enable filtering options in Perseus. Proteins were filtered to retain only those with at least three valid values across all replicates (n=4). Group-wise averaging was applied, requiring a minimum of two valid values out of four replicates, using the mean as the aggregation method.

Data visualization and curve fitting were carried out in Python 3. Proteins were annotated with Gene Ontology (GO) Cellular Compartment. For each protein, the relative isotope abundance (RIA) was calculated using the formula:

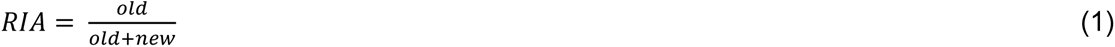

old = intensity of protein synthesized prior to the isotope pulse

new = intensity of protein synthesized post-pulse

This allowed estimation of the fractional replacement of proteins over time. To assess protein turnover, RIA values were plotted against time points (2 h, 8 h, and 24 h) and fitted with the first-order exponential decay model, where *k* is the turnover rate and *x* is the time (2). To calculate the protein half-life (T1/2), the turnover rate *k* was divided by the natural logarithm (3):

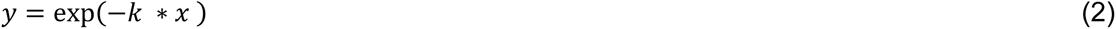

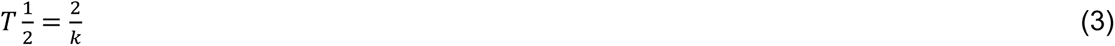

### Statistical analysis

Statistical analyses were performed using GraphPad Prism software (version 9.5.1) and Perseus (version 2.0.9.0). Unless stated otherwise, all data were derived from independent experiments or from independent differentiation experiments for hiPSC-derived neurons. The number of biological samples and independent experimental replicates, from which the data were generated, are provided in the corresponding figure legends. Unless stated otherwise, a two-tailed Mann-Whitney-U-test or two-tailed unpaired t-test was used for comparisons between two groups. For testing effects of one independent variable, one-way analysis of variance (ANOVA) followed by Tukey’s post hoc multiple comparisons test was performed. Two-way ANOVA followed by Tukey’s post hoc multiple comparisons test was used for testing effects of more than one independent variable. The respective tests are indicated in the figure legends. P values < 0.05 were considered statistically significant (*p < 0.05, **p < 0.01, ***p < 0.001).

## Results

### FICD is primarily expressed in dopaminergic neurons in human and rodent substantia nigra

Previously, FICD was reported to be enriched in rat substantia nigra and hippocampus (13). However, given the cellular heterogeneity of the substantia nigra, the neuronal subtype-specific expression of FICD, particularly in dopaminergic neurons primarily affected in PD, remained unclear. We therefore examined FICD expression in dopaminergic neurons of the substantia nigra from rodents and human *post-mortem* brains using immunohistochemistry. In WT rats, we co-immunostained for FICD and the dopaminergic neuron marker tyrosine hydroxylase (TH) (Fig. 1a). Overall, 18.2% of all DAPI-positive cells were positive for FICD (FICD+/DAPI+(%)), of which 80.5% co-expressed TH (TH+/FICD+(%)). In humans, DAB-Ni immunostaining produced blue FICD labeling distinct from brown neuromelanin (NM), a metabolite pigment physiologically present in human dopaminergic neurons (Fig. 1b). A large fraction of dopaminergic neurons showed FICD immunoreactivity (NM+FICD+, Fig. 1b, arrowhead). Specifically, 73.8% NM+ cells co-expressed FICD (Fig. 1b, FICD+/NM+(%)), and 90.1% of FICD+ cells were NM+ (Fig. 1b, NM+/FICD+(%)). The expression of FICD in dopaminergic neurons was independently confirmed by co-immunofluorescence staining for FICD and TH (Fig. S1a). Overall, these data indicate a preferential FICD expression in dopaminergic neurons of both rodent and human substantia nigra.

**Figure 1.**
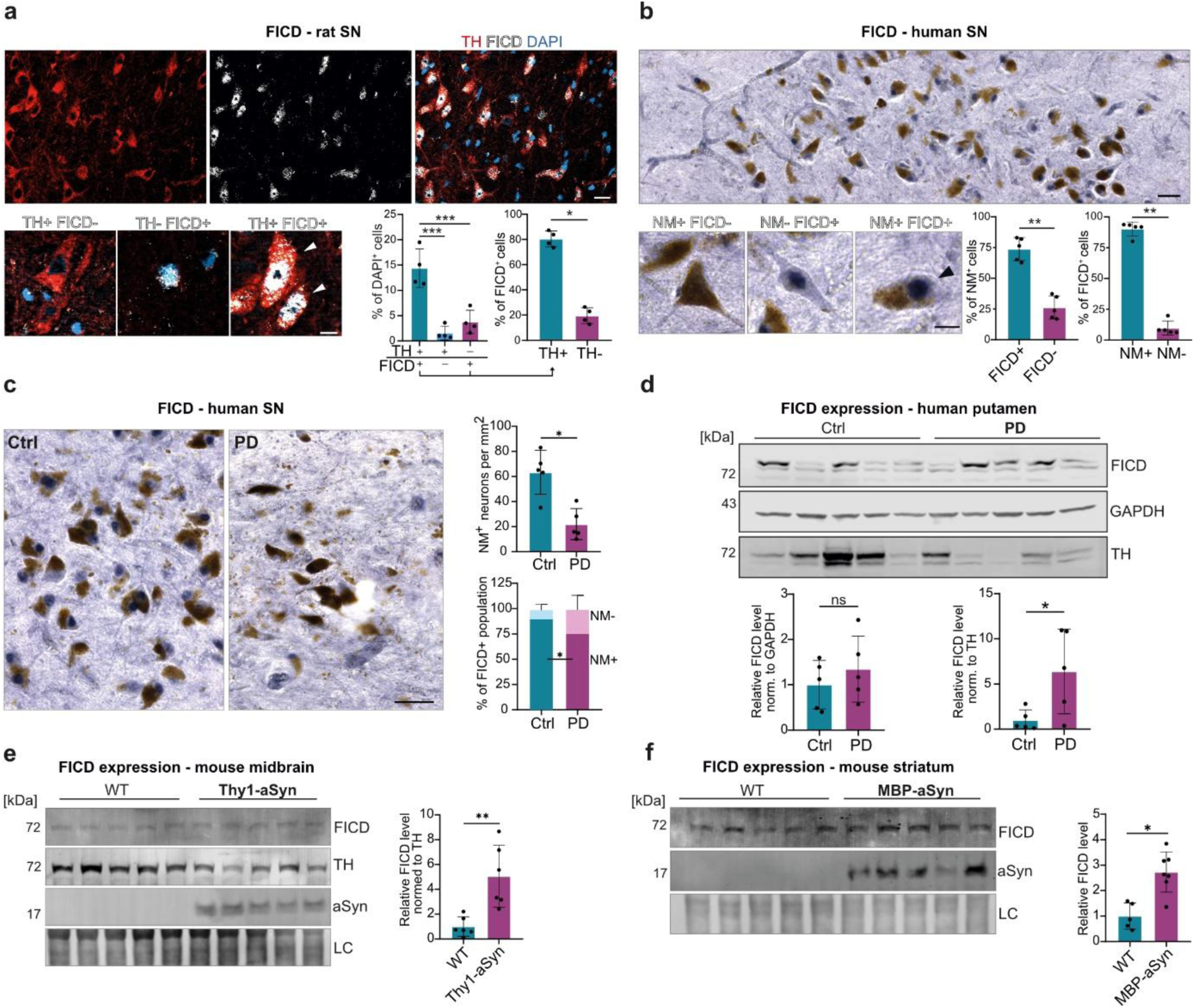
FICD is enriched in dopaminergic neurons and dysregulated in PD and synucleiopathy mice. **a** Immunofluorescence staining of tyrosine hydroxylase (TH, red) and FICD (white), as well as DAPI (blue) in rat substantia nigra. High-magnification example images display TH+FICD-, TH-FICD+, and TH+FICD+ (arrowheads) cells. Percentages of TH+FICD-, TH-FICD+, and TH+FICD+ subpopulations are presented, the unshown TH-FICD- cells comprise the remainder of the population. FICD is preferentially expressed in TH+ dopaminergic neurons (n = 4 animals / group). Scale bar: 5 µm or 20 µm (high-magnification). Statistical analysis: one-way ANOVA with Tukeýs multiple comparisons test and two-tailed Mann-Whitney-U-test. **b** DAB-Ni immunostaining of human substantia nigra from control individuals (Table S1), showing FICD expression (blue) and neuromelanin (NM) pigment (brown), a prototypical feature of dopaminergic neurons. FICD is strongly expressed in NM+ dopaminergic neurons. High-magnification example images display NM+FICD-, NM-FICD+, and NM+FICD+ (arrowhead) cells (n = 5 individuals / group). Scale bar: 50 µm or 20 µm (high-magnification). **c** DAB-Ni immunostaining of FICD in the human *post-mortem* substantia nigra of controls and PD patients. Quantification of NM+ and FICD+ cells demonstrates a significant reduction of NM+ neurons (neurons/mm^2^, right top) and their proportional reduction within FICD+ cells (right bottom) in PD (n = 5 individuals / group). Scale bar: 50 µm. **d** Western blot analysis of human *post-mortem* putamen from controls and PD patients (Table S2) shows no significant difference in FICD levels when normalized to GAPDH (bottom left). However, normalization to TH (bottom right) demonstrates significantly increased FICD levels in PD patients (n = 5 individuals / group). **e-f** Western blot analysis of midbrain tissue from Thy1-aSyn and non-transgenic WT mice. FICD levels were significantly elevated in Thy1-aSyn mice compared with WT controls when normalized to TH (n = 6 animals / group for quantification). **d** Western blot analysis of striatal tissue from MBP-aSyn and non-transgenic WT mice. FICD expression was significantly increased in MBP-aSyn mice compared with WT controls (n= 5-7 animals / group for quantification). Statistical analysis from b - f: two-tailed Mann-Whitney-U-test. p* < 0.05, p** < 0.01, p*** < 0.001. Bar graphs: mean ± SD. *LC: loading control using revert 520 total protein stain; SN: substantia nigra*.

### FICD levels are elevated in the brains of PD patients and respective synucleinopathy mouse models

To evaluate the pathological relevance of FICD in PD, we compared its expression in the *post-mortem* tissue from PD patients with that of control individuals without neurodegenerative disorders. Given the preferential expression of FICD in dopaminergic neurons of the substantia nigra (Fig. 1b) and pronounced dopaminergic neuronal loss in this region (Fig. 1c, right top, neuromelanin staining; Fig. S1b, TH staining), we first performed immunohistochemical analyses and observed a reduced proportion of dopaminergic cells within the FICD expressing population in PD (NM+/FICD+(%): 78.8% in PD versus 90.1% in controls) (Fig. 1c, right bottom). Conversely, the relative proportion of non-dopaminergic FICD-expressing cells (NM-/FICD+(%)) increased.

Next, we performed Western blot analysis of FICD in *post-mortem* putamen, the target region of nigrostriatal dopaminergic projections. When normalized to GAPDH, no significant difference in FICD levels was observed between PD patients and controls (Fig. 1d, bottom left). In contrast, normalization to TH, which was severely reduced in PD, revealed significantly higher FICD levels in patients (Fig. 1d, bottom right), with a trend toward an inverse correlation between FICD and TH levels in PD putamen (Fig. S1c).

We further analyzed FICD expression in two different transgenic mouse models of synucleinopathies, Thy1-aSyn and MBP-aSyn, which overexpress human WT aSyn under the Thy1 and MBP promoter, respectively (26, 27). Thy1-aSyn mice recapitulate the key PD-like phenotypes through neuronal aSyn overexpression, whereas MBP-aSyn mice, which overexpress aSyn in oligodendrocytes, model multiple system atrophy (MSA), an atypical parkinsonian disorder and a synucleinopathy characterized by predominant aSyn aggregation in oligodendrocytes. Consistent with our human data, FICD levels were significantly increased in the midbrain of Thy1-aSyn mice when normalized to TH (given dopaminergic neuronal loss in this model) (Fig. 1e), and were elevated in the striatum of MBP-aSyn mice (Fig. 1f).

Taken together, these converging data from PD patients and corresponding synucleinopathy mouse models demonstrate that FICD is upregulated in association with aSyn pathology and dopaminergic neurons.

### PD patient-derived midbrain neurons carrying an *SNCA* duplication exhibit a selective loss of FICD-expressing dopaminergic neurons, increased AMPylation, and elevated ER-stress

Since the primary characterized function of FICD in humans is the regulation of AMPylation (9), we sought to elucidate the mechanistic link between FICD-mediated AMPylation and aSyn pathology in PD using a human neuronal model. To this end, we employed hiPSC-derived midbrain neurons from a monogenic PD patient carrying an aSyn gene (*SNCA)* duplication (SNCA^Dupl^) and the CRISPR-Cas9-corrected isogenic control (SNCA^Ctrl^) (22). Midbrain neurons were generated from hiPSCs via NPCs (Fig. 2a). As reported in our previous studies, SNCA^Dupl^ neurons displayed increased aSyn aggregation and neuritic phenotypes, including reduced fiber density, as well as neurite volume and diameter (22, 23).

**Figure 2.**
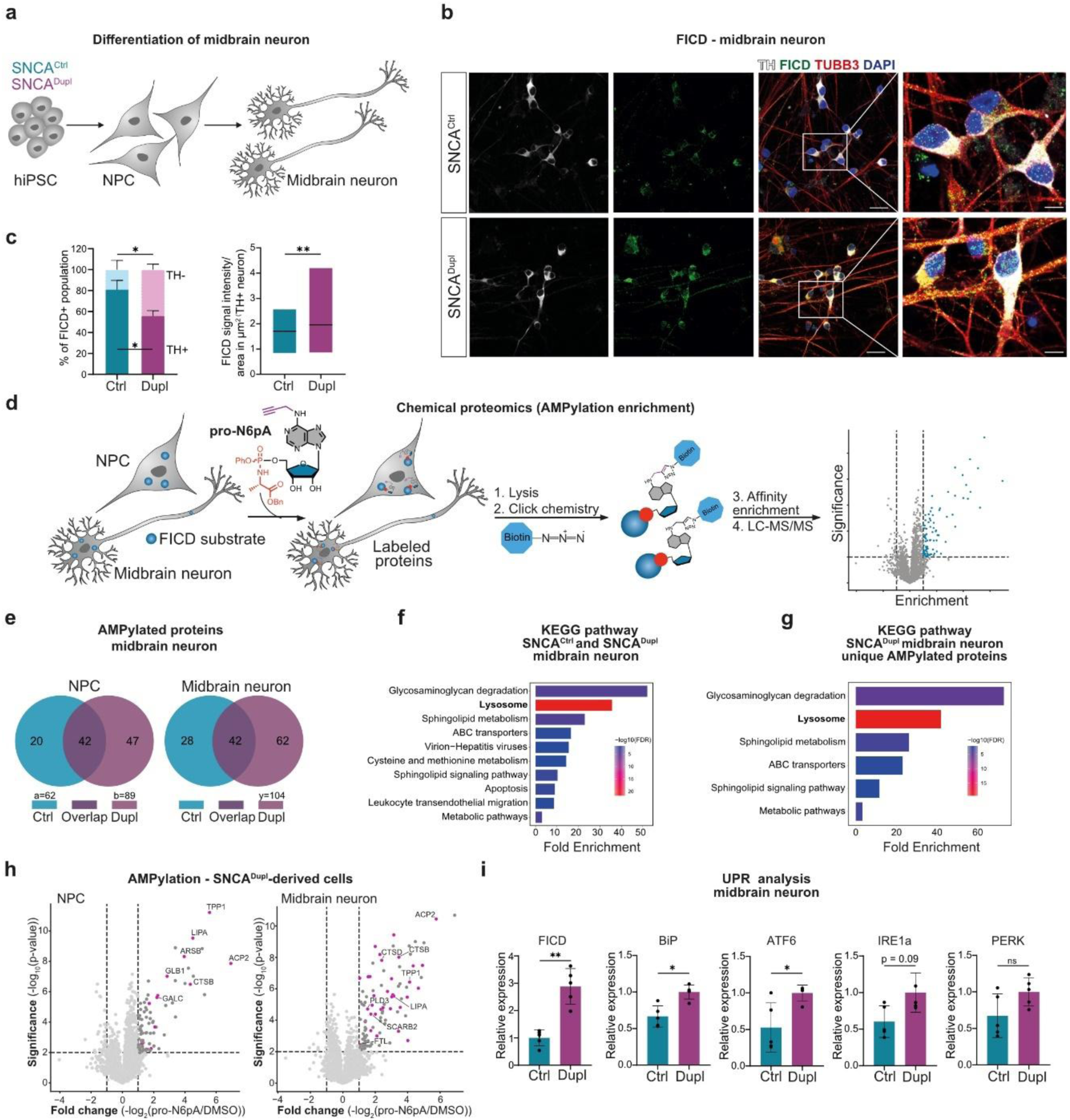
SNCA^Dupl^ midbrain neurons show loss of FICD-positive dopaminergic neurons, with enhanced AMPylation and UPR activation. **a** Schematic hiPSC differentiation workflow from hiPSCs via NPCs to midbrain neurons. **b** Representative triple immunocytochemical images of SNCA^Ctrl^ and SNCA^Dupl^ midbrain neurons stained for TH (white), FICD (green), the pan-neuronal marker TUBB3 (red), and nuclei with DAPI (blue). Scale bars: 20 µm or 5 µm (at higher-magnification). **c** Quantification of TH+ (dark colors) and TH- (light colors) fractions within the FICD+ neuronal population. The proportion of TH+ neurons decreased within the FICD+ population in SNCA^Dupl^-derived midbrain neurons (left); FICD signal intensity shows a significant increase selectively in TH+ SNCA^Dupl^ dopaminergic neurons (right) (n = 3; left: n = 100 individual neurons / group / differentiation; right: n = 64-84 neurons / group /differentiation). Statistical analysis: two-way ANOVA with Tukeýs multiple comparisons test (left), two-tailed Mann-Whitney-U-test (right). **d** Workflow for AMPylation enrichment and detection via chemical proteomics. Intracellular FICD substrates were labelled with the adenosine analog pro-N6pA, followed by biotin labelling using copper-catalyzed click chemistry, affinity enrichment, and LC–MS/MS to enrich probe-labelled proteins. **e** Venn diagram showing the numbers of enriched AMPylated protein hits in NPCs (left) and midbrain neurons (right). **f-g** KEGG pathway enrichment analysis of AMPylated proteins in SNCA^Ctrl^ and SNCA^Dupl^ midbrain neurons (f) or uniquely AMPylated proteins in SNCA^Dupl^ midbrain neurons (g). Bars show fold enrichment; -log10 FDR values are color coded. The “lysosome” pathway shows the strongest statistical significance. **h** AMPylation enrichment in SNCA^Dupl^-derived NPCs (left) and midbrain neurons (right). Volcano plots depict log2 fold change (pro-N6pA vs. DMSO) versus –log10 p-value; dashed lines mark significance cut-offs (p < 0.01, -log2(pro-N6pA/DMSO) > 1). Significantly enriched proteins are shown in dark grey, with lysosomal proteins highlighted in pink. Representative lysosomal proteins are annotated (n = 2; n = 3 technical replicates / experiment). **i** mRNA expression analysis of canonical UPR markers reveals ER-stress phenotype in SNCA^Dupl^ midbrain neurons (n = 5). Bar graphs: mean ± SD. Statistical analysis: two-tailed Mann-Whitney-U-test. p* < 0.05, p** < 0.01. *Ctrl: SNCA^Ctrl^; Dupl: SNCA^Dupl^; TUBB3: βIII-tubulin*.

Consistent with our findings in rodent and human *post-mortem* brains (Fig. 1b-c), immunocytochemical analyses of FICD together with TH, the neuronal marker βIII-tubulin (TUBB3), and DAPI revealed enrichment of FICD in dopaminergic neurons (Fig. 2b). Specifically, 60% SNCA^Ctrl^-derived neurons co-expressed TH and FICD (FICD+TH+/TUBB3+DAPI+(%)), corresponding to over 82% of TH+ neurons being FICD+ (FICD+/TH+(%), Fig. S2a). Notably, the fraction of TH+ dopaminergic neurons within the FICD+ population (TH+/FICD+(%)) was significantly reduced in SNCA^Dupl^ (55.6%) compared with SNCA^Ctrl^ neurons (80.9%) (Fig. 2c, left), while FICD signal intensity was elevated specifically in TH+ neurons (Fig. 2c, right). Thus, SNCA^Dupl^ neurons showed a pronounced loss of FICD-expressing dopaminergic neurons but a higher FICD level in dopaminergic neurons, mirroring the cellular phenotype observed in PD *post-mortem* substantia nigra (Fig. 1c).

To address whether the SNCA^Dupl^-mediated increase in FICD is linked to altered AMPylation activity, we analyzed global protein AMPylation profiles using adenosine analog pro-N6pA-based chemical proteomics (Fig. 2d). AMPylation profiling revealed extensive remodeling of the AMPylated proteome in SNCA^Dupl^ cells (Fig. 2e). We identified 62 significantly enriched AMPylated proteins in SNCA^Ctrl^ NPCs and 89 hits in SNCA^Dupl^ NPCs, whereas 70 versus 104 proteins were identified in differentiated neurons. Overall, PD SNCA^Dupl^ patient-derived cells, in particular the differentiated neurons, exhibited a markedly higher number of uniquely AMPylated proteins than SNCA^Ctrl^ cells (NPCs: 47 versus 20; Neurons: 62 versus 28).

Unbiased KEGG pathway enrichment analysis of all AMPylated proteins identified from both SNCA^Ctrl^ and SNCA^Dupl^ midbrain neurons revealed the lysosomal pathway as the most significantly enriched among the top 10 pathways (Fig. 2f). Importantly, proteins uniquely AMPylated in SNCA^Dupl^ neurons were also most significantly enriched in the lysosomal pathway (Fig. 2g). Notably, AMPylation of lysosome-associated proteins was more extensive and significant in neurons than in NPCs, in which metabolic pathways and amino acid biosynthesis appeared as the most significantly enriched pathways (Fig. 2f-g for neurons, Fig. S2b for respective NPCs).

Among AMPylated lysosome-associated proteins, including CTSB, ACP2, TPP1, and LIPA, were commonly AMPylated, regardless of genotype (SNCA^Dupl^, SNCA^Ctrl^) or differentiation stage (NPCs, neurons) (Fig. 2h for SNCA^Dupl^ cells and Fig. S2c for SNCA^Ctrl^ cells). In contrast, certain AMPylated, lysosome-associated proteins, which have been linked to neurodegeneration, such as CTSD (35), FTL (36), and PLD3 (20, 37) were uniquely AMPylated in SNCA^Dupl^ midbrain neurons (Fig. 2h). Notably, although aSyn was previously reported as a substrate of FICD-mediated AMPylation in cell-free assays (13), it was not detected among enriched AMPylated proteins in both NPCs and midbrain neurons.

FICD is localized to the ER, the primary site of protein AMPylation and of the UPR required to maintain protein homeostasis (38). Given that aSyn pathology impairs UPR responses, and induces ER-stress, as reported in *SNCA* triplication patient-derived neurons (39, 40), the pronounced changes in AMPylation profiles observed in SNCA^Dupl^ neurons raise the question of whether altered AMPylation is associated with UPR dysregulation. We therefore examined the gene expression of FICD and key ER-resident components of the UPR pathway, including BiP, ATF6, IRE1α, and PERK. RT-qPCR analysis demonstrated significant upregulation of FICD, BiP, and ATF6 in SNCA^Dupl^ neurons compared to SNCA^Ctrl^ neurons, suggesting an activated ER-stress in these neurons (Fig. 2i).

Together, PD-derived SNCA^Dupl^ midbrain neurons exhibited a proportional reduction in FICD-expressing dopaminergic neurons, selective upregulation of FICD expression within dopaminergic neurons, increased AMPylation of lysosome-associated proteins, and upregulated canonical UPR markers related to ER-stress. Additionally, a stronger involvement of lysosomal protein AMPylation was observed in differentiated neurons compared with autologous proliferating NPCs.

### Hyperactivation of FICD-mediated AMPylation targets lysosomal proteins and reshapes ER protein processing

Our collective findings from *post-mortem* PD brains, SNCA^Dupl^-derived midbrain neurons, and synucleinopathy mouse models suggest a link between aSyn pathology with activated FICD-AMPylation pathway and enhanced ER-stress. Given the dual enzymatic activity of FICD, which mediates both AMPylation and de-AMPylation, we next asked whether FICD-driven AMPylation contributes causally to these effects.

First, we attempted to overexpress the constitutively AMPylation-active FICD-E234G mutant (9) in SNCA^Dupl^ midbrain neurons by lentiviral transduction of NPCs followed by neuronal differentiation (Fig. S3a). To distinguish the effects of AMPylation from de-AMPylation, NPCs were additionally transduced with lentiviruses encoding FICD-WT or the constitutively inactive FICD-H363A variant (9). Lentiviruses were produced in HEK 293T cells, however, viral particles production of FICD-E234G was markedly reduced compared to FICD-WT and FICD-H363A (Fig. S3b), indicating impaired viral production associated with the active FICD-E234G variant. Subsequent transduction of the FICD variants in NPCs using lentiviral particles at identical multiplicities of infection followed by neuronal differentiation resulted in robust expression of FICD-WT and FICD-H363A in differentiated neurons, whereas FICD-E234G expression was not detectable (Fig. S3c-d). This failure likely reflects detrimental effects of FICD-E234G during viral transfection and integration, and/or cytotoxicity associated with FICD-E234G expression during neuronal differentiation.

To enable controlled modulation of AMPylation activity of FICD in a more tractable system, we next employed plasmid-based overexpression of FICD variants in H4 neuroglioma cells (Fig. 3a). Cells were transfected with FICD-WT, -E234G, -H363A, or an empty vector (mock), which served as a baseline condition, as endogenous FICD expression in H4 cells was barely detectable by Western blot (Fig. S3e). To investigate the interplay between the FICD-AMPylation pathway and aSyn, we utilized H4 cells stably overexpressing human aSyn (H4-aSyn) and corresponding H4 control cells (H4-Ctrl) (Fig. 3a), the latter generated using the identical lentiviral backbone, but lacking aSyn (21, 23).

**Figure 3.**
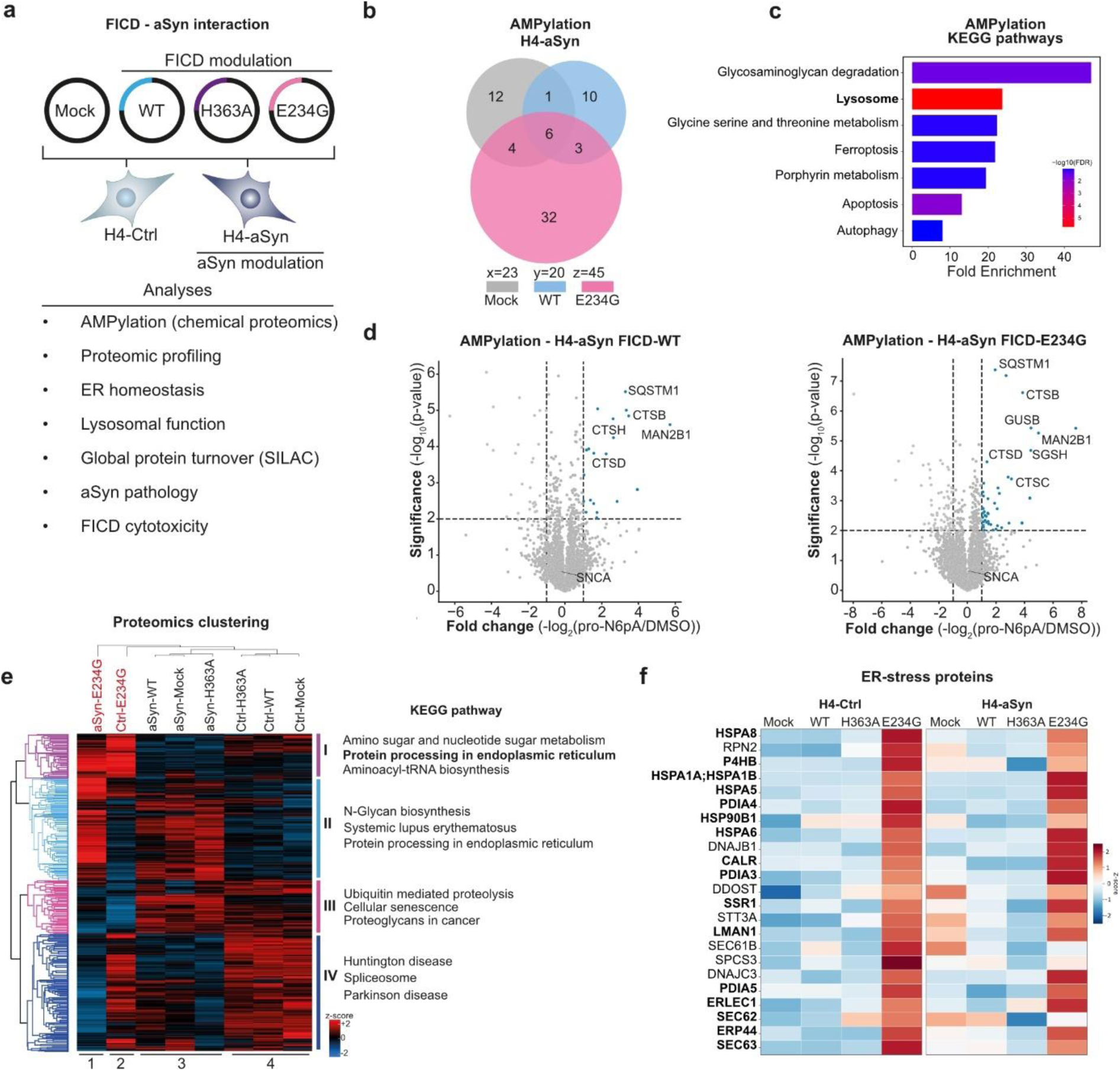
FICD-E234G overexpression elevates AMPylation and reshapes ER-lysosomal pathways. **a** Experimental setup for investigating FICD and aSyn interaction in H4-Ctrl and H4-aSyn cells transiently transfected with FICD-WT, catalytically inactive FICD-H363A, or constitutively active FICD-E234G. Mock transfection served as baseline. **b**. Venn diagram of AMPylated proteins detected in H4-aSyn cells with mock, FICD-WT or FICD-E234G transfection. Overexpression of FICD-E234G resulted in an increase in AMPylated proteins. **c** KEGG pathway enrichment of AMPylated proteins in FICD-WT and FICD-E234G-transfected H4-aSyn cells. The “lysosome” pathway reaches the highest statistical significance. Bar color indicates statistical significance expressed as -log10(FDR). **d** AMPylation enrichment in H4-aSyn cells overexpressing FICD-WT (left) and FICD-E234G (right). Volcano plot depicts log2 fold change (pro-N6pA versus DMSO) versus –log10 p-value; dashed lines mark significance cut-offs (p < 0.01, -log2(pro-N6pA/DMSO) > 1). Significantly enriched AMPylated proteins are highlighted in blue. Representative lysosomal proteins and aSyn (SNCA) are annotated (for c - d: n = 2; n = 2 technical replicates / experiment). **e** Whole proteomic analysis and hierarchical clustering of proteome profiles of H4-Ctrl and H4-aSyn cells, either transfected with mock, FICD-WT, FICD-H363A, or FICD-E234G. Columns represent experimental conditions, partitioned into four clusters (1–4), while rows correspond to differentially regulated proteins (clusters I-IV). On the right, top 3 over-represented KEGG pathways for each row cluster are shown. Z-score: normalized expression scale from lower (blue) to higher (red) levels. The “protein processing in endoplasmic reticulum” pathway is overrepresented in row clusters I and II. **f** Heatmap showing expression of a subset of 23 proteins within the KEEG “protein processing in endoplasmic reticulum” pathway that are upregulated upon FICD-E234G overexpression (For subset identification, refer to Fig. S3i, cluster 5). Proteins involved in the UPR are highlighted in bold (for e - f: n = 3).

To validate the AMPylation-activating effect of FICD-E234G, we performed pro-N6pA-based chemical proteomics using H4-aSyn cells. The number of AMPylated proteins approximately doubled upon FICD-E234G expression, with 45 AMPylated proteins identified, compared with 23 and 20 AMPylated proteins in cells expressing FICD-WT or mock-transfected controls, respectively (Fig. 3b). Specifically, BiP, the well-established FICD target, showed increased AMPylation upon FICD-E234G overexpression (Fig. S3f), as independently confirmed by Western blot analysis (Fig. S3g). Collectively, these results validated robust AMPylation activation by FICD-E234G in the H4-model system.

In accordance with our findings in hiPSC-derived cells (Fig. 2f-g), KEGG pathway analysis of AMPylated proteins again revealed the lysosomal pathway as particularly prominent, showing the highest statistical significance (Fig. 3c). Several lysosomal proteins, including CTSB, CTSD, MAN2B1, and SGSH, were commonly AMPylated in both H4 cells (Fig. 3d) and hiPSC-derived neurons (Fig. 2h). Of note, among these proteins, CTSB exhibited significantly increased AMPylation in H4-aSyn overexpressing FICD-E234G cells compared to FICD-WT cells (Fig. S3f). Consistent with the finding in hiPSC-derived cells (Fig. 2h), aSyn itself was not enriched among AMPylated proteins in H4-aSyn cells. This was further supported by additional analyses including click chemistry, Phos-tag gel, and intact protein mass spectrometry, none of which provided evidence for aSyn AMPylation (Fig. S3i-k).

To define cellular pathways altered by activation of FICD-mediated AMPylation in the context of aSyn overexpression, we performed proteome analysis in cells co-modulated for aSyn and FICD variant expression. Hierarchical clustering was conducted based on protein expression profiles, with experimental conditions arranged in columns and proteins in rows (Fig. 3e). Column clustering showed that mock-, FICD-WT-, and FICD-H363A-transfected cells primarily segregated according to aSyn overexpression (clusters 3 and 4), indicating that aSyn overexpression was the dominant driver of proteomic differences under these FICD conditions. In contrast, H4-Ctrl and H4-aSyn cells expressing FICD-E234G formed separate clusters (clusters 1 and 2), indicating that hyperactivation of FICD-mediated AMPylation induces additional proteomic alterations beyond those driven by aSyn alone. Moreover, row clustering identified four major protein clusters (I-IV). In order to identify molecular pathways affected through the interplay between FICD and aSyn, KEGG pathway analysis of proteins from individual row clusters was performed. This revealed enrichment of protein homeostasis-related pathways, including “Protein processing in the endoplasmic reticulum” (within clusters I-II), “Ubiquitin mediated proteolysis” (within cluster III), as well as neurodegenerative disorder-related pathways, such as “Huntington disease” and “Parkinson disease” (within cluster IV).

Given FICD’s prominent role within the ER, we performed an in-depth analysis of 112 proteins identified within the “Protein processing in the endoplasmic reticulum” pathway. Hierarchal clustering revealed a subset of 23 proteins (Fig. S3k) that were consistently upregulated in both H4-Ctrl and H4-aSyn cells upon FICD-E234G expression. Notably, UPR-related proteins, such as BiP (HSPA5), were upregulated within this subset (Fig. 3f).

In summary, unbiased chemical proteomics and global proteomics demonstrate that hyperactivation of FICD-mediated AMPylation broadly increases cellular AMPylation, preferentially targeting lysosomal proteins, and profoundly reshapes the ER protein processing machinery.

### Hyperactivation of FICD-mediated AMPylation leads to ER-stress and decreased CTSB activity

We next investigated the functional consequences of AMPylation hyperactivation, focusing on protein homeostasis pathways. Due to increased ER-stress markers in SNCA^Dupl^ neurons (Fig. 2i) and upregulation of UPR-related proteins upon specific AMPylation hyperactivation (Fig. 3e), we systematically examined key UPR signaling branches as crucial indicators of ER-stress. In addition, given the prominent AMPylation of lysosomal proteins in both hiPSC-derived neurons (Fig. 2f-g) and H4 cells (Fig. 3c) we assessed lysosomal function.

The UPR restores protein homeostasis under ER-stress via three sensor branches (Fig. 4a): IRE1, PERK, and ATF6. IRE1 mediates XBP1 mRNA splicing (sXBP1), PERK phosphorylates eIF2α to induce CHOP expression, and ATF6 translocates to the Golgi to form an active transcription factor. BiP, regulated by FICD-mediated AMPylation (41), coordinates these responses (42). RT-qPCR analysis revealed increased BiP gene expression upon FICD-E234G overexpression in both H4-Ctrl and H4-aSyn cells. Further, we observed an IRE1- and PERK- mediated stress response, evident by increased expression of spliced XBP1P downstream of IRE1, and elevated expression of PERK and CHOP in the PERK branch (Fig 4b). Notably, increased ER-stress appears to be a common effect of hyperactivation of FICD-mediated AMPylation in both H4-Ctrl and H4-aSyn cells.

**Figure 4.**
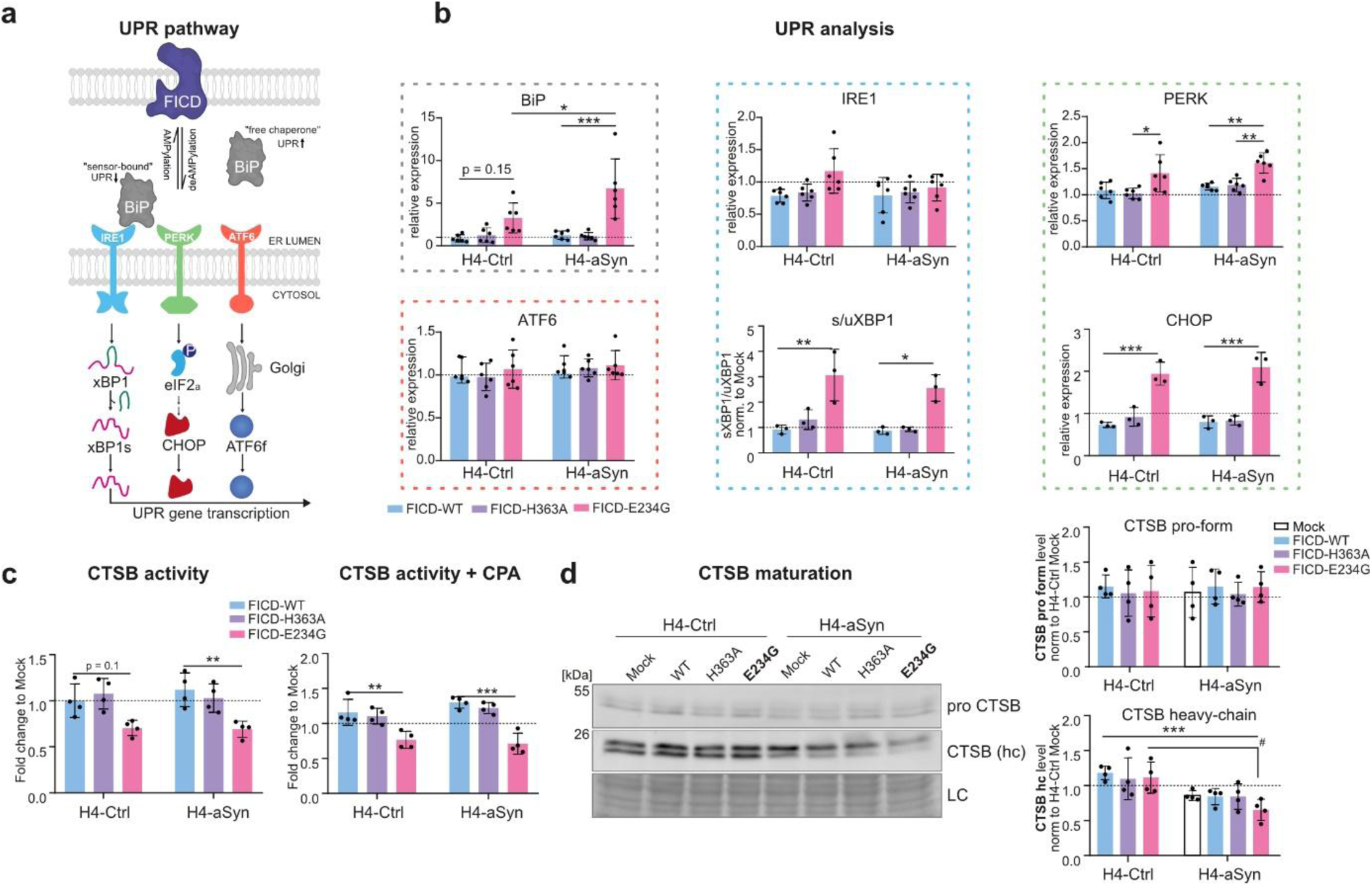
Hyperactivation of FICD-mediated AMpylation induces ER-stress and impairs lysosomal protease activity. **a** Schematic representation of the effect of FICD on BiP AMPylation with the downstream regulatory impact on UPR signaling branches IRE1α, PERK, and ATF6. **b** RT-qPCR analysis of canonical UPR markers in H4-Ctrl and H4-aSyn cells transfected with mock, FICD-WT, FICD-H363A, or FICD-E234G. mRNA expression of BiP, spliced XBP1 in the IRE1 branch, as well as PERK and CHOP in the PERK branch, is significantly upregulated upon FICD-E234G expression. Data are shown as expression relative to mock (dashed line) (n = 3 – 6). **c** CTSB activity assay normalized to mock (dashed line). Left, H4-Ctrl and H4-aSyn cells modulated by FICD variants. Right, FICD-modulated cells additionally exposed to cyclopiazonic acid (CPA; ER Ca^2+^-ATPase inhibitor) (n = 3). **d** Western blot showing CTSB pro-form (right top) and cleaved, mature CTSB heavy-chain (right bottom) in H4-Ctrl and H4-aSyn cells transfected with mock, FICD-WT, FICD-H363A, or FICD-E234G (n = 4). Bar graphs: mean ± SD. Statistical analysis: two-way ANOVA with Tukeýs multiple comparisons test. p* < 0.05, p** < 0.01, p*** < 0.001. *LC: loading control via Coomassie brilliant blue; hc: heavy chain. sXBP1: spliced variant of XBP1; uXBP1: unspliced XBP1*.

Since CTSB and CTSD are important lysosomal enzymes (43) and among the frequently AMPylated substrates in SNCA^Dupl^-derived neurons (Fig. 2h) and H4-aSyn cells (Fig. 3d), both models exhibit aSyn dyshomeostasis (22, 23), we examined their activity under AMPylation hyperactivation. We observed significantly reduced CTSB activity in H4-aSyn cells overexpressing FICD-E234G (Fig. 4c, left). This effect was further exacerbated by additional treatment with cyclopiazonic acid (CPA), a Ca^2+^-ATPase inhibitor that depletes ER calcium stores and induces ER-stress (Fig. 4c, right). Although the synergistic effect of AMPylation activation and CPA was evident in both H4-Ctrl and H4-aSyn cells, the response was more pronounced in H4-aSyn cells. Since functional CTSB requires post-translational maturation during trafficking along the ER-Golgi-endosome-lysosome pathway, including PTMs and proteolytic processing, we performed Western blot analysis to assess CTSB cleavage. While pro-CTSB levels were not significantly affected by FICD modulation within H4-Ctrl or H4-aSyn cells (Fig. 4d), the enzymatically active CTSB heavy chain was significantly reduced in H4-aSyn cells overexpressing FICD-E234G, indicating impaired CTSB maturation. In contrast to CTSB, CTSD activity was not significantly affected by AMPylation activation (Fig. S4), suggesting a preferential targeting of CTSB by dysregulated FICD activity.

In summary, hyperactivation of FICD-mediated AMPylation increases ER-stress responses and impairs CTSB activity and maturation, with the effects on CTSB being particularly prominent under aSyn overexpression.

### Hyperactivation of FICD-mediated AMPylation decreases overall protein turnover rate

Building on our finding that specific hyperactivation of FICD-mediated AMPylation disrupts ER homeostasis (Fig 3e-f, 4b), preferentially AMPylates lysosomal substrates (Fig. 3d), and impairs CTSB activity (Fig. 4c-d), we hypothesized that this combined phenotype would disrupt overall protein turnover. To assess proteome-wide protein turnover, we performed a pulse-chase experiment in cell culture using stable isotope labeling with amino acids in cell culture (SILAC), followed by proteomic analysis (Fig. 5a). After confirming successful isotope labeling (Fig. S5a), we detected total 2450 proteins with a mean half-life of 15.79 h (Fig. S5b). Of these, 938 proteins were consistently detected under all conditions and included in a hierarchical clustering analysis (Fig. 5b). In both H4-Ctrl and H4-aSyn cells, mock and FICD-WT transfected cells clustered closely (clusters 2 and 4), while FICD-E234G transfected cells formed distinct clusters (clusters 1 and 3), in line with the whole-proteome finding that AMPylation activation via FICD-E234G overexpression drives large differences in protein landscapes (Fig. 3e). To gain further insight, we analyzed protein half-lives by grouping the SILAC-identified proteins according to their subcellular compartments or structures (Fig. 5c). FICD-E234G transfection increased global protein half-lives in both H4-Ctrl and H4-aSyn cells. Comparison between both cell lines showed no major differences, except for actin cytoskeleton proteins, which exhibited significantly decreased half-lives in H4-aSyn cells as compared to H4-Ctrl cells (Fig. S5c).

**Figure 5.**
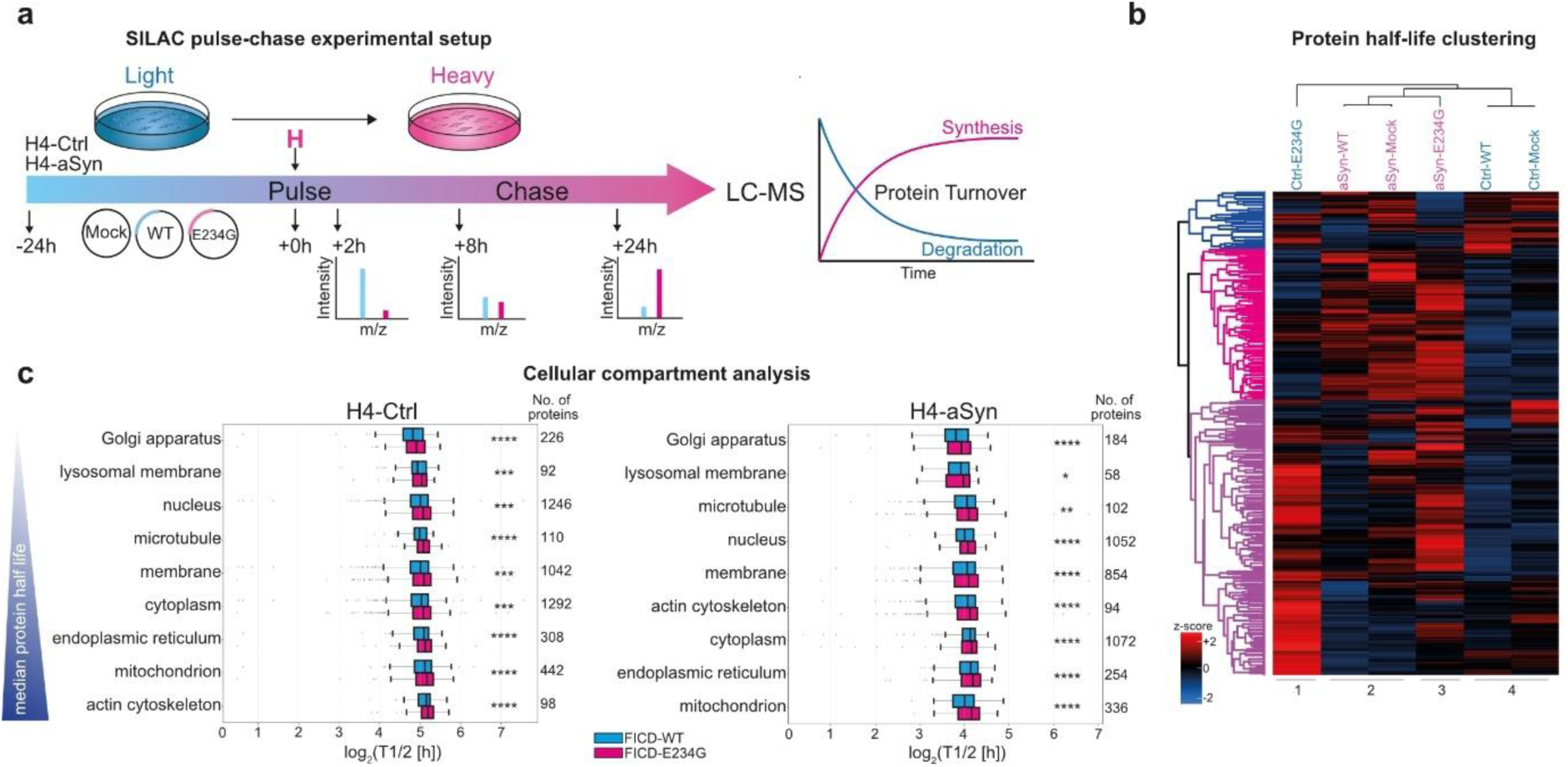
SILAC proteomics reveals global reduction in protein turnover upon hyperactivation of FICD-mediated AMPylation. **a** Schematic overview of the SILAC pulse-chase experimental setup. H4-Ctrl and H4-aSyn cells were initially cultured in SILAC media containing light or heavy isotopes. Twenty-four hours after transfection with mock, FICD-WT, or FICD-E234G, the pulse-chase was initiated by switching the SILAC media to the opposite isotope condition (light-to-heavy or heavy-to-light). Isotope incorporation was monitored at multiple chase time points (2 h, 8 h, and 24 h) by LC-MS/MS to quantify the overall protein turnover. Depicted is the pulse-chase experimental workflow initiated with light SILAC medium. **b** Hierarchical clustering of half-lives from 938 commonly identified proteins. Z-score: normalized half-lives scale from lower (blue) to higher (red) levels. **c** Subcellular compartment analysis. Boxplots show distribution of protein half-lives grouped by GO cellular compartment annotation in H4-Ctrl (left) and H4-aSyn (right) cells. The x-axis shows the log2-transformed protein half-lives in log_2_(T1/2) [h]. Each box spans the interquartile range (IQR: 25^th^-75^th^ percentile), with the center line indicating the median. Whiskers extend to data points within 1.5× IQR from the lower and upper quartiles, outliers are represented by dots. FICD-E234G overexpression significantly reduces global protein turnover compared with FICD-WT overexpression in both H4-Ctrl and H4-aSyn cells (n = 4). Statistical analysis: two-tailed Wilcoxon-matched-pairs signed rank test. p* < 0.05, p** < 0.01, p*** < 0.001, p**** < 0.0001.

### Hyperactivation of FICD-mediated AMPylation increases aSyn aggregation and induces apoptosis

Given the regulatory role of FICD in protein homeostasis, we investigated whether it modulates aSyn aggregation, a crucial pathogenic mechanism of PD. We employed two complementary assays for aSyn aggregation: a solubility assay to quantify insoluble aSyn and a filter trap assay to detect aggregated aSyn species. Analyses were performed in H4-aSyn cells, as H4-Ctrl cells express barely detectable levels of aSyn (Fig. S6a). We observed significantly increased levels of insoluble aSyn in FICD-E234G transfected H4-aSyn cells compared to mock-, FICD-WT-, or FICD-H363A-transfected cells (Fig. 6a). Concordantly, the filter trap-based aggregation assay confirmed elevated levels of aSyn aggregates in FICD-E234G overexpressing cells (Fig. 6b), an effect that was particularly pronounced when using the aggregates-specific antibody MJFR-14-6-4-2, as previously established (44).

**Figure 6.**
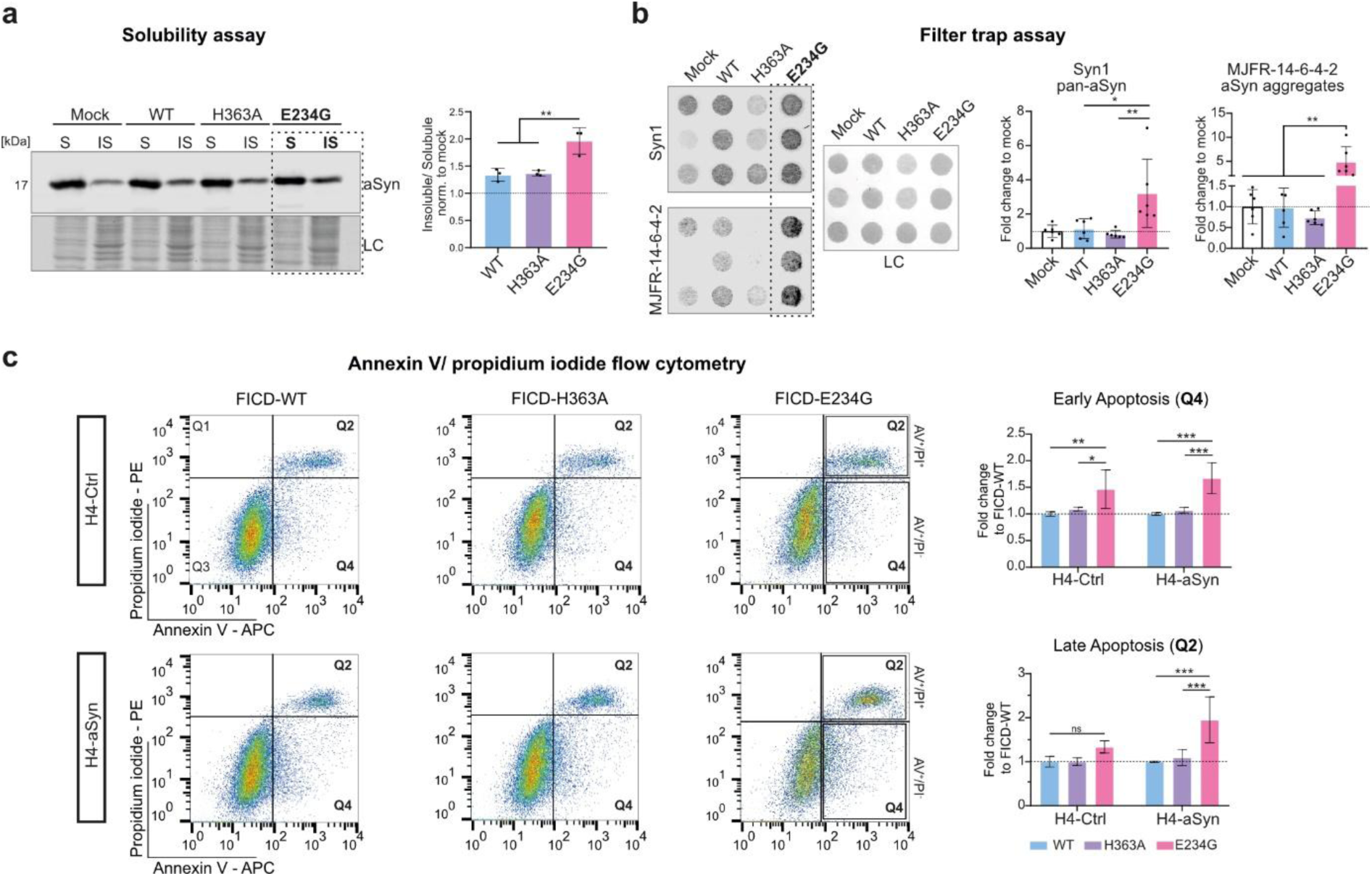
Hyperactivation of FICD-mediated AMPylation promotes aSyn aggregation and induces apoptosis in H4-aSyn cells. **a** Solubility assay of aSyn in H4-aSyn cells transfected with FICD-WT, -H363A, or -E234G using Western blot, analyzing aSyn in soluble (S) and insoluble (IS) fractions. Quantification shows elevated levels of insoluble aSyn upon FICD hyperactivation (n = 3). **b** Filter trap assay detecting aSyn aggregates using a pan-aSyn antibody (Syn1) and a conformation-dependent antibody with a higher affinity to aSyn aggregates (MJFR-14-6-4-2). Quantification demonstrates an increase in aSyn aggregation upon expression of FICD-E234G compared to FICD-WT or FICD-H363A (n = 6). Statistical analysis for a and b: one-way ANOVA with Tukeýs multiple comparisons test. **c** Flow cytometric analysis of apoptosis in H4-aSyn and H4-Ctrl cells. Cells were stained with Annexin V and propidium iodide (PI) to distinguish early (Q4) and late (Q2) apoptotic populations (highlighted in the left panel). Expression of FICD-E234G significantly increased early apoptosis in both cell lines, while late apoptosis was selectively elevated in H4-aSyn cells (n = 3). Statistical analysis: two-way ANOVA with Tukeýs multiple comparisons test. Bar graphs: mean ± SD. p* < 0.05, p** < 0.01, p*** < 0.001. Dashed lines: mock controls. *LC: total protein via Coomassie brilliant blue for a) and Ponceau S for b); AV: Annexin V; PI: propidium iodide; IS: insoluble fraction; S: soluble fraction*.

We next assessed potential toxicity associated with hyperactivation of FICD-mediated AMPylation by measuring apoptosis using flow cytometry (Fig. 6c). Cells were stained with Annexin V and propidium iodide (PI) to identify early (Annexin V+/PI-) and late apoptotic (Annexin V+/PI+) populations (Fig. 6c and Fig. S6b for gating setup). Expression of the active FICD-E234G variant significantly increased early apoptosis in both H4-Ctrl and H4-aSyn cell lines (Fig. 6c). In contrast, late apoptosis was significantly induced only in H4-aSyn cells, suggesting a pronounced cytotoxic response to FICD hyperactivation in the presence of aSyn aggregates.

### Inhibition of FICD-mediated AMPylation by closantel ameliorates aSyn aggregation and neuritic phenotype in SNCA^Dupl^ midbrain neurons

Efforts to identify small-molecule pharmaceutical modulators interfering with the FICD-AMPylation pathway have intensified in recent years (45–47). Fluorescence polarization assays using fluorescent ATP analogs and recombinant FICD have enabled the discovery of pharmaceutical inhibitors of FICD-mediated AMPylation, including closantel, a halogenated salicylanilide (47). Given the increased aSyn aggregation and neuritic defects previously observed in SNCA^Dupl^ midbrain neurons (22, 23), we tested whether closantel treatment has the potential to mitigate these phenotypes. Inhibition efficacy was assessed by Western blot analysis of BiP AMPylation as a surrogate marker. We confirmed a significant reduction in the AMPylated-to-total BiP ratio upon closantel treatment in SNCA^Dupl^ neurons (Fig. 7a, Fig. S7a), without detectable cytotoxicity (Fig. S7b). In contrast, closantel did not alter BiP AMPylation in SNCA^Ctrl^ neurons under identical treatment conditions (Fig. 7a). A similar lack of AMPylation inhibition effect was observed in H4 cells expressing distinct FICD variants, despite baseline differences between FICD variants (Fig. S7c-d). These results indicate that closantel selectively inhibits AMPylation both in a cell type-dependent manner and in the presence aSyn pathology.

**Figure 7.**
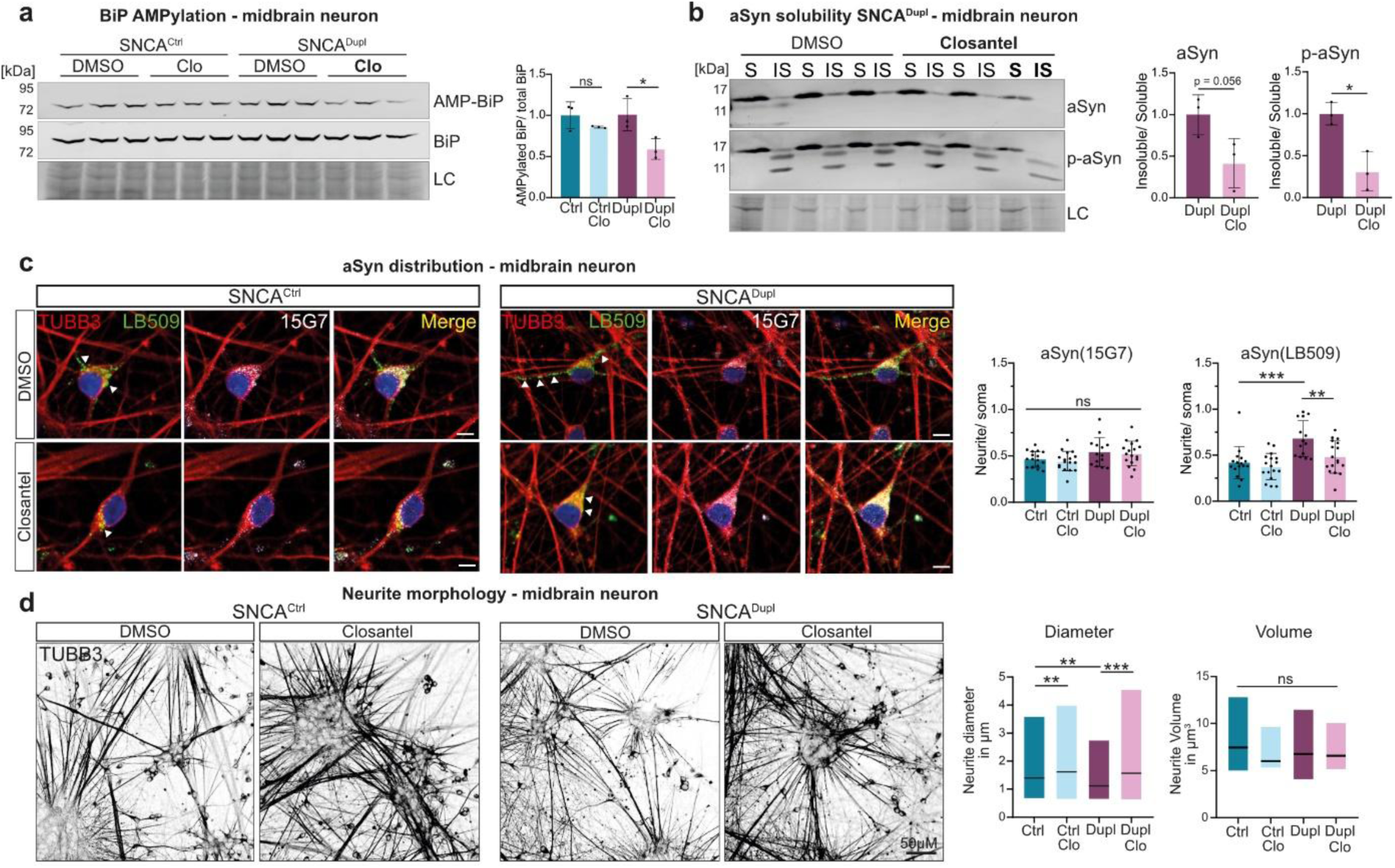
AMPylation inhibition by closantel rescues SNCA^Dupl^-induced aSyn aggregation and neuritic morphology in midbrain neurons. **a** Western blot analysis of AMPylated and total BiP in SNCA^Ctrl^ and SNCA^Dupl^ midbrain neurons treated with closantel or DMSO as control. Quantification of the AMPylated-to-total BiP ratio revealed a significant reduction upon closantel treatment in SNCA^Dupl^ midbrain neurons, confirming effective AMPylation inhibition (n = 3). Statistical analysis: one-way ANOVA with Tukeýs multiple comparisons test. **b** Solubility assay of aSyn and phosphorylated aSyn in SNCA^Dupl^ neurons treated with closantel or DMSO. Following closantel treatment, levels of insoluble total aSyn were reduced (p = 0.056). Importantly, insoluble phosphorylated aSyn was significantly and more robust reduced (n = 3). Statistical analysis: two-tailed unpaired t test. **c** Immunocytochemical analysis of aSyn using antibodies 15G7 (white) and LB509 (green, higher affinity to aggregated aSyn) in SNCA^Ctrl^ and SNCA^Dupl^ midbrain neurons expressing TUBB3 (red). Nuclei were stained via DAPI (blue). LB509-immunopositive aSyn (highlighted by arrows) accumulated predominantly in neurites of SNCA^Dupl^ neurons, whereas closantel treatment significantly reduced aSyn accumulation in neurites. No effect was observed in SNCA^Ctrl^ neurons. Quantifications represent the ratio of neuritic to somatic fluorescence intensity (n=3; n = 16, 17, 14, 17 neurons for Ctrl, Ctrl-Clo, Dupl and Dupl-Clo, respectively). Statistical analysis: one-way ANOVA with Tukeýs multiple comparisons test. Bar graphs for a-c: mean ± SD. **d** Analysis of neuritic morphology in SNCA^Ctrl^ and SNCA^Dupl^ neurons treated with closantel. Left: representative immunostaining and maximum intensity projection images of neurons labelled by TUBB3 immunostaining (black); Right: quantification shows an increased neurite diameter in both genotypes following treatment, while neurite volume remained unchanged (n = 3; for diameter: n = 210, 174, 236, 161 neurites for Ctrl, Ctrl-Clo, Dupl and Dupl-Clo conditions, respectively; for volume: n = 20 neurites / differentiation). Scale bars: 50 µm. Statistical analysis: one-way ANOVA with Tukeýs multiple comparisons test. Floating bars: min to max, line at media. p* < 0.05, p** < 0.01, p*** < 0.001. *AMP-BiP: AMPylated BiP; Ctrl: SNCA^Ctrl^; Dupl: SNCA^Dupl^; Clo: closantel; IS: insoluble fraction; LC: loading control using revert 520 total protein stain; p-aSyn: phosphorylated aSyn; S: soluble fraction; TUBB3: βIII-tubulin*.

To explore potential effects of closantel on aSyn aggregation, we performed solubility assay followed by Western blot analysis using a pan-aSyn antibody in soluble and insoluble protein fractions. We observed a trend toward decreased solubility of aSyn in SNCA^Dupl^ hiPSC-derived midbrain neurons after closantel treatment, although this change did not reach statistical significance (Fig. 7b, aSyn). In contrast, analysis of phosphorylated aSyn at serine 129, a PTM that is markedly elevated in Lewy body-associated aSyn (48), revealed a significant reduction in the ratio of insoluble versus soluble aSyn carrying this PTM (Fig. 7b, p-aSyn). Building on our previous observation of preferential aSyn accumulation in neurites of SNCA^Dupl^ neurons (22), we analyzed the subcellular distribution of total and aggregated aSyn in soma and neurites using immunocytochemistry. We employed a pan-aSyn antibody (15G7) and an antibody (LB509) with a higher affinity for aggregated aSyn (Fig. 7c). Both antibodies have been established for detecting differential accumulation of aSyn species along neurites in these cells (22). While the subcellular distribution of total aSyn (15G7) remained unchanged, aggregated aSyn (LB509) accumulated predominantly in neurites of SNCA^Dupl^ neurons, consistent with our prior findings (22). Importantly, closantel treatment significantly reduced the accumulation of aggregated aSyn in neurites. Notably, closantel did not affect either aSyn solubility (Fig. S7e) or its neuritic accumulation in SNCA^Ctrl^ neurons (Fig. 7c).

Because SNCA^Dupl^ neurons show neuritic defects, we next examined neuritic diameter and volume following closantel treatment. Closantel increased neuritic diameter in SNCA^Dupl^ neurons (Fig. 7d), restoring partially the neuritic phenotypes that are characteristic of these cells (22, 23).

Together, pharmacological inhibition of the FICD-AMPylation pathway by closantel reduced BiP AMPylation and selectively alleviated aSyn aggregation in SNCA^Dupl^ midbrain neurons. In addition, closantel ameliorated aSyn-mediated neuritic deficits in these neurons by increasing neurite diameter.

## Discussion

AMPylation is the best-characterized function of FICD (9, 49). Our findings provide mechanistic insights into the role of dysregulated FICD activity in PD, thereby highlighting its potential as an interventional target for PD.

We demonstrate the pathological relevance of FICD dysregulation in PD-associated neurodegeneration. Extending a previous observation of predominant FICD expression in rat substantia nigra (13), we show that FICD is preferentially expressed in dopaminergic neurons in both rodent and human substantia nigra, as well as in hiPSC-derived midbrain neurons. In PD substantia nigra and SNCA^Dupl^ PD patient-derived midbrain neurons, the proportion of dopaminergic FICD+ cells was reduced, whereas the fraction of non-dopaminergic FICD+ neurons increased proportionally. Consistently, FICD and TH levels were inversely correlated in PD brains. Moreover, FICD expression was elevated in brains from PD patients and synucleinopathy mouse models, as well as in PD-derived dopaminergic neurons, thereby linking increased FICD levels to reduced dopaminergic integrity.

Multiple lines of evidence from this study demonstrate a bidirectional interaction between aSyn pathology and the FICD-mediated AMPylation pathway, providing mechanistic insight into the pathogenic contribution of this interplay in PD. On the one hand, aSyn pathology alters the FICD-AMPylation pathway and related proteostatic mechanisms: specifically, aSyn overexpression induces upregulation of FICD in SNCA^Dupl^-derived NPCs and neurons, as well as in mouse brains; SNCA^Dupl^ promotes AMPylation, as supported by increased global AMPylation activity in SNCA^Dupl^ NPCs and neurons; SNCA^Dupl^ results in a selective loss of dopaminergic FICD-expressing neurons. Moreover, in aSyn-overexpressing H4 cells (H4-aSyn), additional FICD hyperactivation reduced CTSB activity and triggered apoptosis. Given that aggregated aSyn disrupts ER integrity and protein-folding capacity (39), the observed aSyn-mediated effects can be explained by exacerbated aSyn pathology, which may interfere with FICD function in the ER and downstream protein trafficking, either directly or indirectly. On the other hand, dysregulation of the FICD-AMPylation pathway influences aSyn aggregation. In H4-aSyn cells, we provide direct evidence that forced activation of FICD-mediated AMPylation indeed promotes aSyn aggregation, likely via AMPylation-driven lysosomal dysfunction, consistent with observations in *C. elegans* (*12*). This is further supported by the co-upregulation of the FICD-AMPylation pathway and aSyn aggregation in SNCA^Dupl^ neurons. Together, our data suggest a self-amplifying feedback loop between aSyn pathology and FICD-mediated AMPylation, in which FICD deAMPylation activity may be counteracted, thereby affecting UPR and the ER-lysosome signaling, particularly in such susceptible dopaminergic midbrain neurons, as previously shown for other long-lived cells (50).

Our results highlight a functional crosstalk between FICD-mediated AMPylation and protein homeostasis pathways. High-throughput AMPylation profiling across diverse cell models revealed a recurring lysosomal signature, indicating that AMPylation of lysosomal targets modulates their processing, subcellular localization, and consequently function, despite ER localization of FICD. This suggests a crucial compartmental crosstalk along the ER-Golgi-endosome-lysosome axis. Indeed, in H4 cells, increased CTSB AMPylation via FICD hyperactivation correlated with reduced CTSB activity, consistent with our previous cell-free assays (6). Similarly, dysregulated AMPylation of PLD3, a key lysosomal single-strand DNA exonuclease and phospholipase (51), impacts its maturation and activity, as reported previously (20, 52). Moreover, we provide insights into a causal link between excessive activation FICD-mediated AMPylation, elevated BiP AMPylation, and ER-stress using the H4 cell model. As the key ER chaperone and gatekeeper of the UPR, BiP is a well-established AMPylation target. AMPylation of BiP inactivates its chaperone function by reducing affinity for unfolded proteins and facilitates its dissociation from UPR sensors. Under homeostatic conditions, AMPylation of BiP serves to remove excess BiP (53, 54), however, hyperactivation of BiP AMPylation depletes the pool of active BiP chaperons, thereby triggering chronic UPR signaling and ER-stress. Beyond direct effects on individual target proteins, our data demonstrate global effects of FICD dysregulation on cellular protein homeostasis. In H4 cells, hyperactivation of FICD-mediated AMPylation led to delayed overall protein turnover, ER-stress, lysosomal dysfunction, and consequently apoptosis, consistent with the observations in SNCA^Dupl^-derived neurons, where increased AMPylation is linked to elevated ER-stress.

Our data additionally revealed distinct AMPylation signatures depending on the neuronal differentiation state. Specifically, differentiated neurons display a broader AMPylation repertoire than proliferating NPCs, while SNCA^Dupl^-derived cells exhibit overall increased AMPylation. These findings suggest that AMPylation is dynamic, finely tuned during neuronal differentiation and remodeled under disease conditions.

It should be noted that our results from H4-aSyn cells and human NPCs and neurons did not detect aSyn AMPylation using complementary approaches, despite a previous report from cell-free assays (13). As aSyn lacks an ER-targeting signal peptide, it is not primarily localized to the ER lumen, likely explaining the absence of AMPylation in the present cell models. Nevertheless, aSyn associates or interacts with the ER, particularly under overexpression, aggregation, or stress conditions (55–57). Thus, aSyn AMPylation cannot be entirely excluded; its undetectable levels may reflect both low abundance and the stringent substrate specificity of FICD within cellular environments. Taken together, our data support that the observed effects of dysregulated FICD-AMPylation pathway occur via an aSyn AMPylation-independent mechanism.

The collective data presented in this study support toxic effects of dysregulated FICD-mediated AMPylation in the context of PD pathology. In H4 cell models, forced hyperactivation of FICD causally promotes apoptosis, indicating that excessive AMPylation is detrimental to cell viability. Consistently, overexpression of the constitutively active FICD-E234G proved challenging in hiPSC-derived neurons, suggesting a heightened vulnerability of neurons to excessive FICD activity. Supporting the neurotoxicity of dysregulated FICD-AMPylation pathway, SNCA^Dupl^ neurons which exhibit markedly elevated AMPylation activity, displayed pronounced neuritic deficits. Importantly, pharmacological inhibition of FICD-mediated AMPylation using closantel reversed multiple SNCA^Dupl^-induced phenotypes, including aSyn aggregation, its accumulation along the neurites, and neuritic deficits, further supporting a causal and pathological role of dysregulated FICD-mediated AMPylation. Taken together, preferential FICD expression in dopaminergic neurons, combined with the detrimental effects of FICD-mediated AMPylation, may underlie the selective loss of FICD-expressing dopaminergic neurons observed in SNCA^Dupl^-derived models and *post-mortem* substantia nigra, supporting a pathological role for dysregulated FICD signaling. The toxic potential of excessive FICD activity for AMPylation is also supported by independent studies: in *Drosophila*, overexpression of FICD-E234G is deleterious, whereas endogenous FICD-WT, with dual AMPylation and de-AMPylation activities, buffers proteotoxic stress (58). Moreover, the homozygous FICD-R371S mutation, which causes infancy-onset diabetes with a severe neurodevelopmental phenotype, impairs BIP deAMPylation and is associated with dysregulated ER-stress signaling (11). Consistent with our findings, a study in *C. elegans* demonstrated that constitutive activation of FICD promotes aSyn aggregation, however, aggregation in worms was associated with improved mobility and survival (12). This discrepancy likely reflects context-dependent effects of aggregation. In *C. elegans,* inclusion formation may be protective by sequestering toxic oligomers, whereas in highly polarized human neurons, aggregates accumulating in neurites can impair axonal transport, disrupt ER-lysosome communication, and exacerbate proteostatic stress. Differences in the extent and regulation of FICD activity across models may further contribute to these divergent outcomes. Nevertheless, these collective findings highlight the therapeutic potential of targeting FICD-mediated AMPylation in PD treatment, while emphasizing the need for precise modulation to avoid adverse effects.

As a potential intervention, we tested closantel as a modulator of FICD-AMPylation and observed beneficial effects in human neurons. Given that two independent synucleinopathy mouse models also exhibited FICD dysregulation in affected brain regions, further studies are needed to validate the therapeutic effects of modulating this pathway *in vivo.* Importantly, while closantel demonstrated restoration of aSyn pathology and neuronal phenotypes, reported adverse effects in humans (59), underscore the need to search for improved mimics. Future work could focus on developing structurally optimized and safer derivatives, potentially offering a promising avenue for therapeutic intervention in PD.

## Conclusion

We provide insights into the pathological relevance of FICD-mediated AMPylation in PD-related neurodegeneration and highlight its contribution to aSyn aggregation through a bidirectional interplay with aSyn pathology. Mechanistically, dysregulated FICD-mediated AMPylation acts as a critical molecular switch for intracellular protein homeostasis. Aberrantly upregulated FICD activity disrupts cellular homeostasis along the ER-lysosome axis, promoting aSyn pathology and cell death. Importantly, pharmacological inhibition of FICD-mediated AMPylation reverses aSyn aggregation and neuritic degeneration, underscoring the potential of targeting this pathway as a therapeutic intervention in PD.

## List of abbreviations

aSyn: alpha-synuclein
BiP: Binding-immunoglobulin protein
CPA: Cyclopiazonic acid
Ctrl: Control
CTSB: Cathepsin B
CTSD: Cathepsin D
Dupl: Duplication
ER: Endoplasmic reticulum
FICD: Filamentation induced by cAMP domain-containing protein
hiPSC: Human-induced pluripotent stem cell
LB: Lewy body
LC-MS: Liquid chromatography mass spectrometry
MS: Mass spectrometry
MSA: Multiple system atrophy
NM: Neuromelanin
NPC: Neural progenitor cell
p-aSyn: phosphorylated alpha-synuclein
PBS: phosphate buffered saline
PD: Parkinsońs disease
PI: Propidium iodide
PBS: phosphate buffered saline
pro-N6pA: N^6^-propargyl adenosine phosphoramidate
PTM: Post-translational modification
PurMA: purmorphamine
RT-qPCR: Reverse transcription quantitative polymerase chain reaction
SILAC: Stable isotope labeling with amino acids in cell culture
UPR: Unfolded protein response
WT: Wild type

## Declarations

### Declarations Ethics approval and consent to participate

All procedures involving the generation and use of hiPSCs, including skin biopsy collection and informed consent were approved by the local Institutional Review Board (Nr. 259_17B) of Friedrich-Alexander-University Erlangen-Nürnberg, Erlangen, Germany. Breeding, housing, and brain preparation of transgenic mice (Thy1-aSyn (Line 61), MBP-aSyn (Line 29)) and their littermates, as well as of Sprague-Dawley WT rats were approved by the local governmental administrations for animal health (TS03-20, Division for Animal Welfare, Friedrich-Alexander-University Erlangen-Nürnberg, Erlangen Germany; AZ. 55.2,2-2532-2-1489, AZ. 55.2-DMS 2532-2-218 Regierung von Unterfranken, Würzburg).

### Author contributions

Conceptualization: AK, JW, PK, WX; Writing – original draft: AK, LH, PK, WX; Writing - review and editing: AK, LH, AB, MA, TGB, GES, SR, JW, PK, WX; Methodology: AK, LH, AB, AS, YS, MA, TB, FZ, SR, PK, WX; Resources: SR, FZ, TBD, GES, JW, PK, WX; Investigation: AK, LH, AS, AB, TB, JW, PK, WX; Data curation: AK, LH, AB, PK, WX; Validation: AK, LH, AS, PK, WX; Visualization: AK, LH, PK, JW, WX; Formal analysis: AK, LH, PK, WX; Software: AK, LH, PK, WX; Supervision: JW, PK, WX; Project administration: PK, WX; Funding acquisition: FZ, TGB, GES, PK, JW, WX.

## Acknowledgements

We thank the donor for providing precious human skin biopsy for hiPSC generation. We thank Holger Meixner, Sonja Plötz, Judith Stemick, Verena Schmidt, Martin Weidenfeller for their excellent technical support

