## Supplementary material for "Aberrant FICD-mediated AMPylation drives α-Synuclein pathology and overall protein dyshomeostasis in dopaminergic neurons in Parkinson’s disease"

**Correspondence:**

Pavel Kielkowski

### Supplementary figures

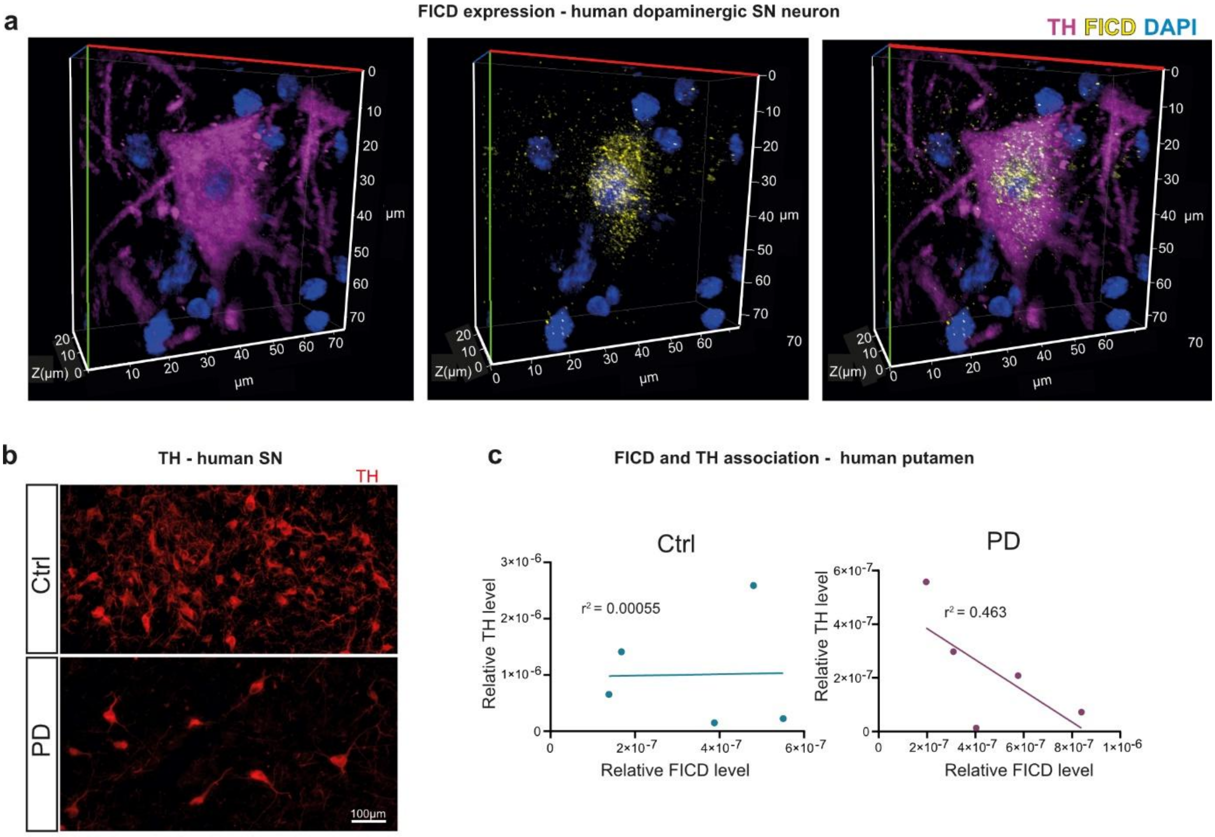


**Figure S1. Analysis of FICD and TH in human *post-mortem* substantia nigra.**

**a** Confocal z-stack (82 optical sections) of human substantia nigra tissue stained for tyrosine hydroxylase (TH, magenta), FICD (yellow), and nuclei (DAPI, blue), visualized as a 3D projection using Zen 2012 (Zeiss). The image illustrates FICD expression within TH-positive dopaminergic neurons. Axes indicate spatial dimensions in µm. **b** Representative immunofluorescence images showing TH signals in the substantia nigra pars compacta of controls and PD patients, illustrating a substantial loss of TH immunoreactivity in PD. **c** Association analysis of TH and FICD protein expression levels based on Western blot analysis of *post-mortem* putaminal tissue from controls and PD patients shown in Figure 1d. Linear regression lines and coefficient of determination $r^2$ values are shown in the scatter blots. No association was observed in controls ($r^2$ = 0.0055), whereas a trend of negative correlation was detected in PD samples ($r^2$ = 0.463, p = 0.21). *SN: substantia nigra.*

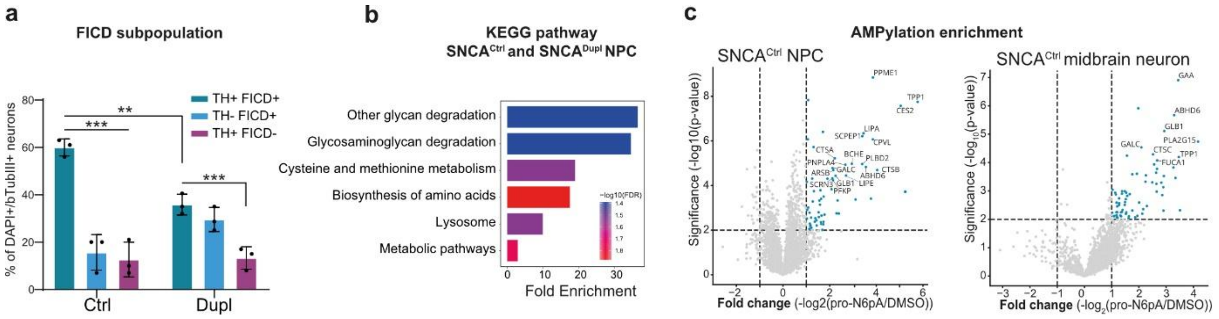


**Figure S2. FICD subpopulation analysis and AMPylation profiling in hiPSC-derived cells.**

**a** FICD subpopulation analysis showing percentage of TH+FICD+, TH-FICD+, and TH+FICD- neurons (βIII-tubulin+) in SNCA$^{Ctrl}$ and SNCA$^{Dupl}$-derived midbrain neurons. The unshown TH-FICD- cells comprise the remainder of the population (n = 3). Bar graphs: mean ± SD. FICD expression in TH+ and TH- neurons was quantified in 100 neurons per *SNCA* genotype in each experiment. Statistical analysis: two-way ANOVA with Tukey´s multiple comparisons test. **b** KEGG pathway enrichment of AMPylated proteins in SNCA$^{Ctrl}$ and SNCA$^{Dupl}$-derived NPCs. Bar color indicates statistical significance expressed as -log10(FDR). **c** AMPylation enrichment in SNCA$^{Ctrl}$ NPCs (left) and midbrain neurons (right). Volcano plots depict log2 fold change (pro-N6pA versus DMSO) versus –log10 p-value; dashed lines mark significance cut-offs (p < 0.01, -log2(pro-N6pA/DMSO) > 1). Significantly enriched AMPylated proteins are highlighted in blue, with example lysosomal proteins annotated. (n = 2; n = 3 technical replicates / experiment).

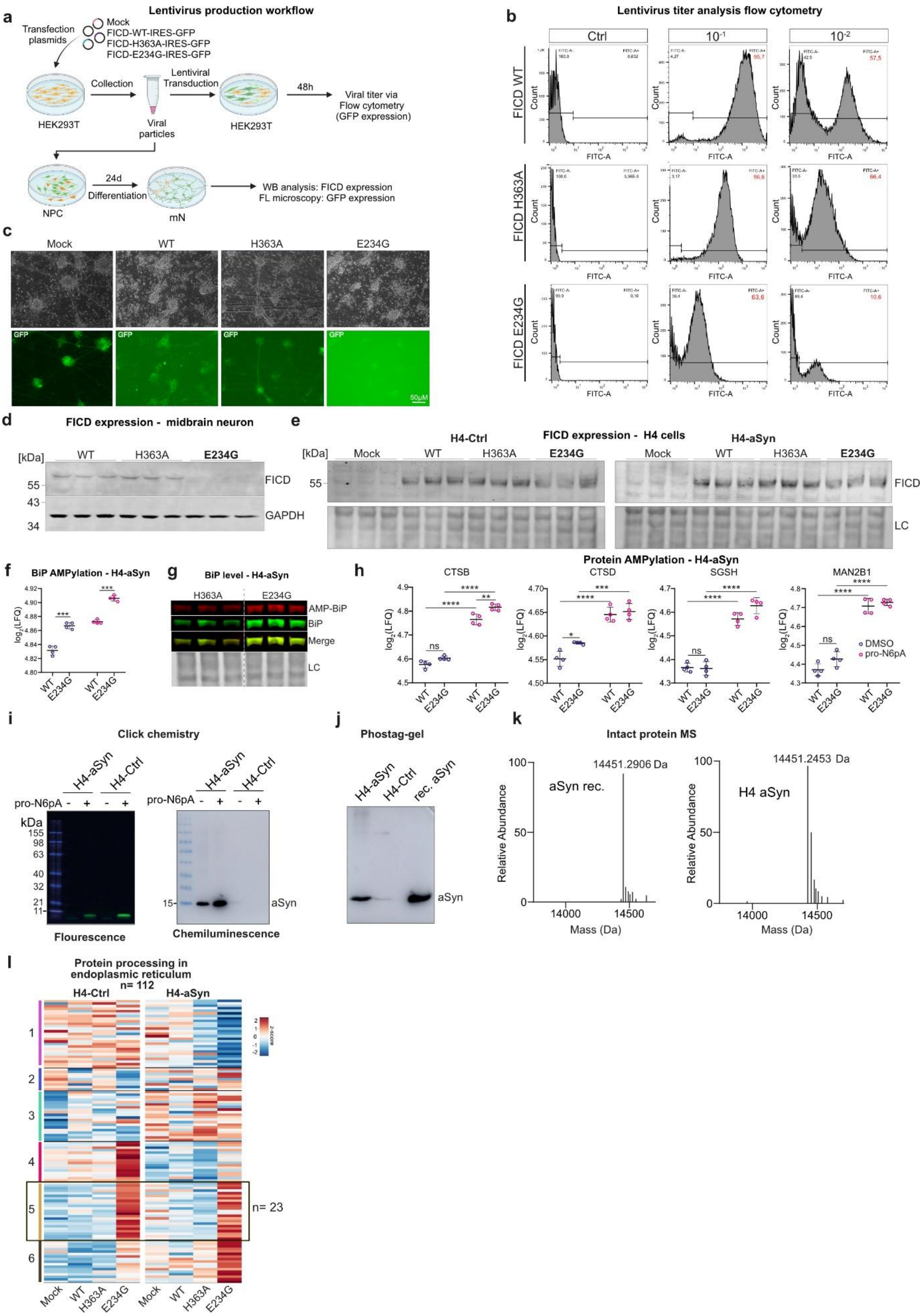


**Figure S3. Lentivirus generation, FICD modulation, AMPylation analysis and proteomics.**

**a** Workflow for lentivirus production and quality control in HEK293T cells with subsequent lentiviral transduction of iPSC-derived NPC and differentiation into midbrain neurons. Virus containing FICD variant-IRES-GFP, allowing co-expression of FICD variants and GFP. Figure created with https://BioRender.com. **b** Flow cytometric analysis of lentiviral titers. HEK293T cells were transduced with serial dilutions of viral particles ($10^{-1}$,1:10; $10^{-2}$,1:100). Ctrl: non-infected cells. Histograms display GFP-fluorescence intensity (FITC-A) and were used to determine transduction efficiency of differently diluted FICD-WT, FICD-H363A and FICD-E234G lentiviruses. **c** Representative phase-contrast (top) and fluorescence (bottom) images of differentiated midbrain neurons transduced with the indicated lentiviral constructs (multiplicity of infection (MOI) = 1). GFP expression is not detectable in FICD-E234G-transduced cells, indicating failed transduction in these cells. Scale bar: 50 µm. **d** Western blot analysis of FICD levels in transduced $SNCA^{Dupl}$ midbrain neurons, demonstrating robust FICD expression in FICD-WT- and FICD-H363A-transduced neurons. In FICD-E234G-transduced neurons, FICD expression is absent. GAPDH was used as a protein loading control. **e** Western blot analysis of FICD levels in H4-Ctrl (left) and H4-aSyn (right) cells following mock or plasmid transfection with FICD-WT, FICD-H363A, or FICD-E234G. In both cell lines, endogenous FICD (mock-transfected cells) were barely detectable. In contrast, transfection with FICD variants resulted in robust overexpression of the respective proteins. **f** Quantification of AMPylated BiP in H4-aSyn cells transfected with FICD-WT or FICD-E234G, analyzed by chemical proteomics, showing a significantly increased BiP AMPylation upon FICD-E234G overexpression (n = 2 independent experiments with n = 2 technical replicates/ experiment). **g** Validation of FICD-E234G-induced BiP AMPylation in H4-aSyn cells by Western blot. AMPylated BiP (detected using an AMP-Thr antibody, red) and total BiP (detected using a BiP antibody, green), together with the merged channels, are shown. FICD-E234G increased AMPylated BiP levels compared to FICD-H363A-overexpressing cells. **h** Quantification of AMPylated lysosomal proteins (CTSB, CTSD, SGSH, MAN2B1) analyzed by chemical proteomics. Data represent $\log_2$ (Label-free quantification (LFQ)) intensities in DMSO- or pro-N6pA-treated H4-aSyn cells transfected with FICD-WT or FICD-E234G (n = 2; n = 2 technical replicates/ experiment).

Statistical analysis: two-way ANOVA with Tukey´s multiple comparisons test. **i** Click chemistry coupled with in-gel fluorescence analysis (left) of aSyn AMPylation status from H4-aSyn cells treated with pro-N6pA probe after ‘click reaction’ with TAMRA-azide as described previously (1). Western blot analysis using chemiluminescence and the aSyn antibody Syn 1 was performed as a control to confirm the position of aSyn (~ 15 kDa) on the blot (right). In-gel fluorescence analysis failed to detect AMPylated aSyn in the probe-treated H4-aSyn cells at the ~ 15kDa (left). **j** Phos-tag gel analysis of aSyn, capable of distinguishing unmodified aSyn from its phosphorylated or AMPylated counterparts, failed to detect any modified aSyn species in H4-aSyn cells. Recombinant human aSyn, generated as described previously (2), was used as a control to visualize unmodified aSyn on Western blot. **k** Intact protein mass spectrometry analysis of aSyn isolated from H4-aSyn cells (right) and recombinant aSyn (left) to analyze aSyn AMPylation status. No significant difference in the apparent molecular mass of aSyn from H4-aSyn cells was observed when compared to human recombinant (rec.) aSyn. **l** Cluster analysis of proteins belonging to the "Protein processing in endoplasmic reticulum" pathway (n = 112) in H4-Ctrl and H4-aSyn cells across Mock, FICD-WT, FICD-H363A, and FICD-E234G conditions. The black box highlights a subset of proteins (cluster 5, n = 23) specifically upregulated in the FICD-E234G condition. Color scale represents z-scored abundance. $p^{**} < 0.01$. *LC, loading control with Revert 520 Total Protein Stain; MS, mass spectrometry. FL, fluorescence*

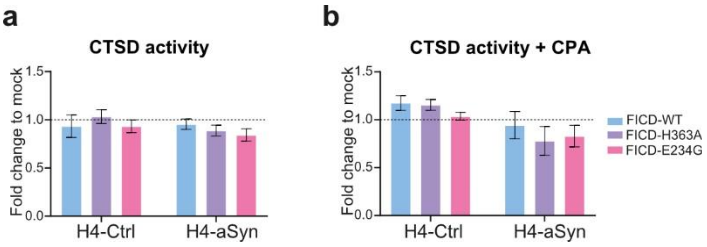


**Figure S4. Cathepsin D activity assay in FICD-modulated H4-Ctrl and H4-aSyn cells.**

**a** Cathepsin D (CTSD) activity assay in H4-Ctrl or H4-aSyn cells transfected with mock, FICD-WT, FICD-H363A, or FICD-E234G. CTSD activity was normalized to mock (dashed line). **b** CTSD activity assay in cells additionally treated with cyclopiazonic acid (CPA) (n = 3). Bar graphs: mean ± SD. Statistical analysis: two-way ANOVA with Tukey´s multiple comparisons test.

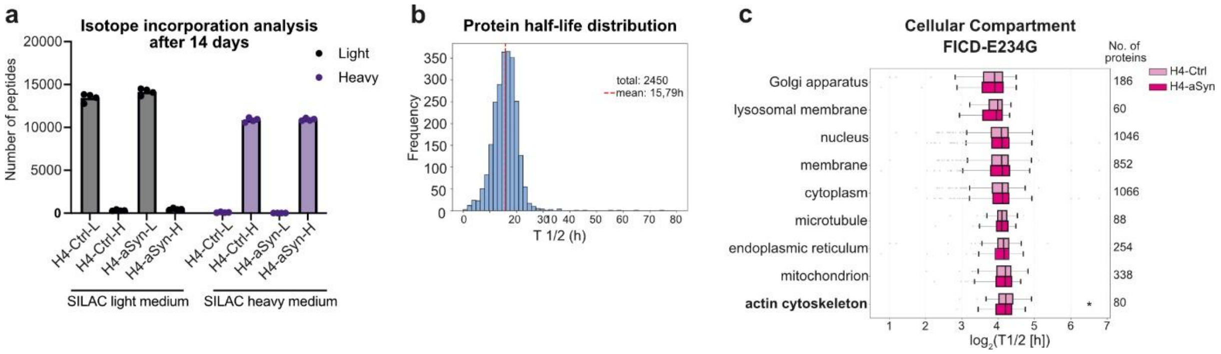


**Figure S5. SILAC pulse-chase proteomics: labeling control and compartment-resolved half-life analysis**.

**a** LC-MS/MS validation of isotope incorporation**.** After 14 days in light (L) or heavy (H) SILAC medium, H4-Ctrl and H4-aSyn cells show a successful isotope incorporation. Depicted are the incorporation efficiencies with SILAC light (left) and heavy (right) media, respectively. **b** Histogram of protein half-lives (T1/2, hours) of all measured proteins (n = 2450, mean half-life = 15.79 h) from SILAC pulse–chase experiments across FICD and aSyn conditions, including mock-, FICD-WT-, and FICD-E234G-transfected H4-aSyn and H4-Ctrl cells. **c** Subcellular compartment analysis. Boxplots show distribution of protein half-lives grouped by GO Cellular Compartment annotation in FICD-E234G-transfected H4-Ctrl and H4-aSyn cells. Proteins involved in actin cytoskeleton only demonstrates significantly slower protein turnover in H4-aSyn cells overexpressing FICD-E234G compared to their H4-Ctrl counterparts. The x-axis shows the log2-transformed protein half-lives in $\log_2$(T1/2) [h]. Each box spans the interquartile range (IQR: 25$^{th}$-75$^{th}$ percentile), with the center line indicating the median. Whiskers extend to data points within 1.5× IQR from the lower and upper quartiles, outliers are represented by dots (n = 4). Statistical analysis: two tailed Wilcoxon-matched-pairs singed rank test. p* < 0.05. *APC: allophycocyanin; PE: phycoerythrin.*

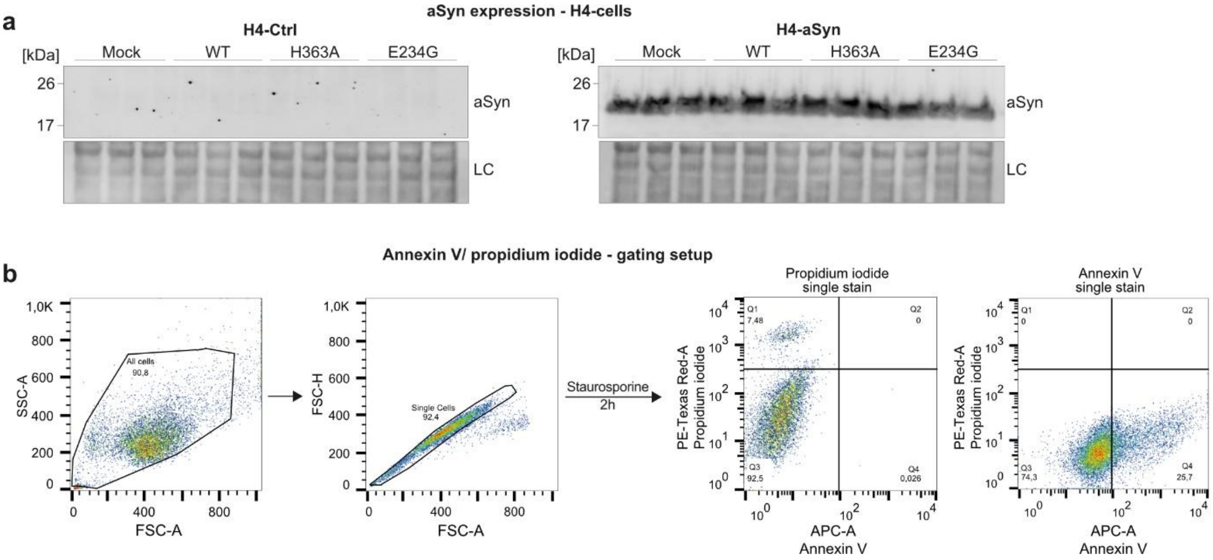


**Figure S6. aSyn expression in H4-Ctrl and H4-aSyn cells and flow cytometric gating strategy for apoptosis assay.**

**a** Western blot analysis of aSyn levels in H4-Ctrl (left) and H4-aSyn (right) cells transfected with mock, FICD-WT, FICD-H363A, and FICD-E234G. aSyn is barely detectable in H4-Ctrl cells. **b** Gating strategy for Annexin V/ propidium iodide apoptosis assay. After treating cells with staurosporine, an inducer of non-lytic apoptosis, single staining for either Annexin V or propidium iodide was used to identify necrotic (PI+, Q1) or apoptotic (Annexin V+, Q4) cell populations. *LC, loading control with Revert 520 Total Protein Stain; AV: Annexin V; PI: propidium iodide*.

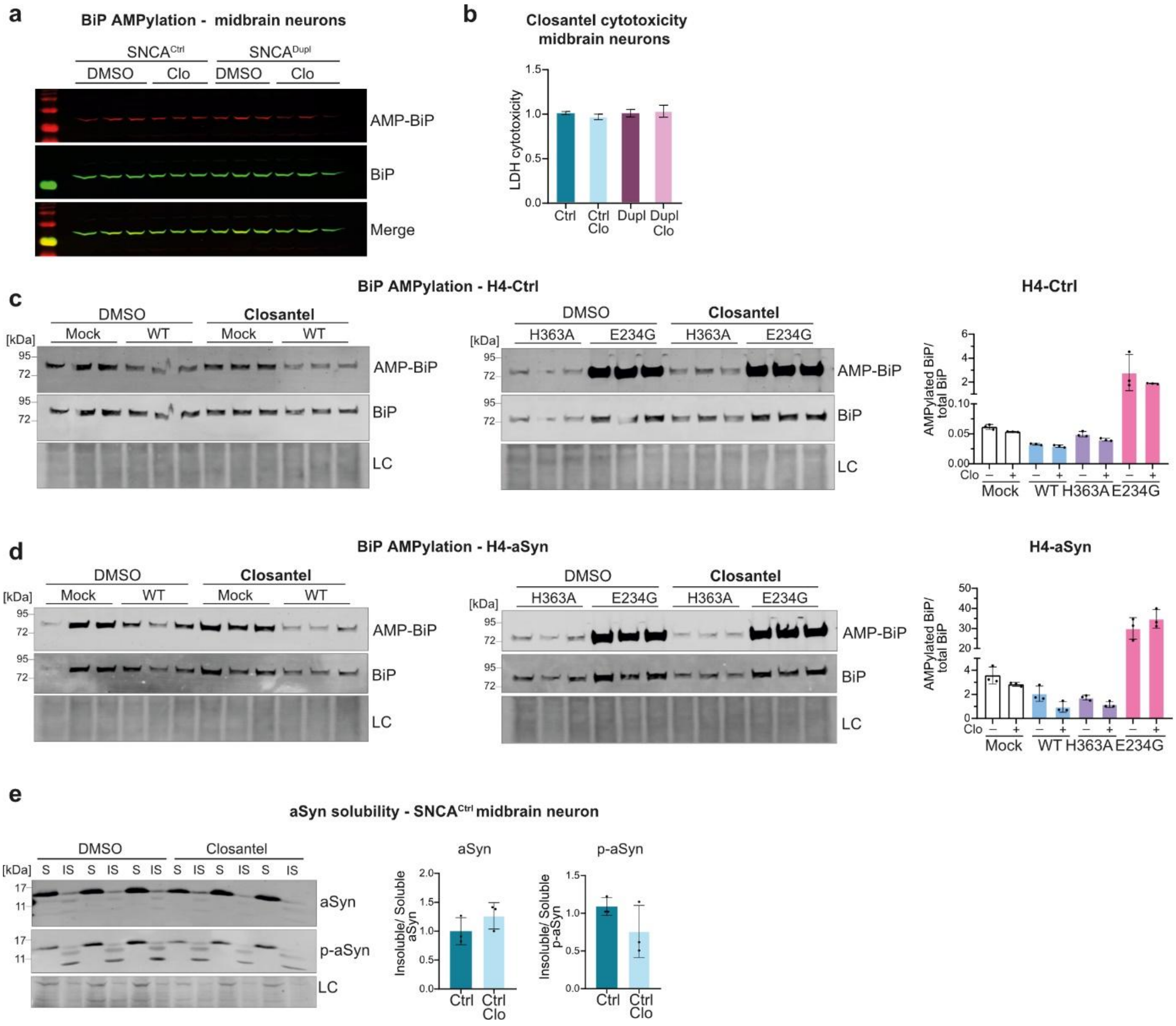


**Figure S7. The effects of closantel on BiP-AMPylation and cytotoxicity in hiPSC-derived neurons, its impact on BiP AMPylation in H4 cells, and its influence on aSyn aggregation in SNCA$^{Ctrl}$ midbrain neurons.**

**a** Western blot of DMSO- and closantel-treated SNCA$^{Ctrl}$ and SNCA$^{Dupl}$ midbrain neurons. Immunostaining of AMPylated BiP and total BiP, shown in Fig. 7a, is presented in red (AMPylated BiP) and green (total BiP), along with the merged signal of both channels. **b** LDH cytotoxicity assay demonstrating no toxic effect in hiPSC-derived midbrain neurons treated with closantel compared to DMSO (n = 3). Bar graphs: mean ± SD. Statistical analysis: two-way ANOVA with Tukey´s multiple comparisons test. **c-d** Western blots and quantification of AMPylated BiP-to-total BiP ratios in H4-Ctrl (c) and H4-aSyn (d) transfected with mock, or FICD variants (WT, H363A, and E234G). Although FICD-E234G increases BiP AMPylation in H4-Ctrl or H4-aSyn as expected, closantel treatment shows no effect on ratios of AMPylated

BiP/total BiP in FICD variant-overexpressing H4-Ctrl (c) or H4-aSyn cells (d) compared to their respective untreated counterparts (n = 3); Bar graphs: mean ± SD. Statistical analysis: ordinary one-way ANOVA with Tukey´s multiple comparisons test. **e** aSyn solubility assay in DMSO- or closantel-treated $SNCA^{Ctrl}$ midbrain neurons. Western blots of soluble and insoluble fractions with pan-aSyn (aSyn) and phosphorylated aSyn (p-aSyn) antibodies. Quantification of ratio of soluble and insoluble aSyn show no significant change following closantel treatment (n = 3). Bar graphs: mean ± SD. Statistical analysis: two-tailed Mann-Whitney-U-test. *AMP-BiP: AMPylated BiP; Ctrl:* $SNCA^{Ctrl}$*; Dupl:* $SNCA^{Dupl}$*; Clo: closantel; IS: insoluble fraction; LC: loading control using revert 520 total protein stain; p-aSyn: phosphorylated aSyn: S: soluble fraction.*

**Table S1.** Demographic and clinical data of *post-mortem* substantia nigra tissue donors (Brain and Body Donation Program of Sun City, Arizona)

| Diagnosis | Sex | Age (yrs) | Post mortem interval (h) | Brain weight (g) | Braak NF score | Braak Lewy Stage |
|---|---|---|---|---|---|---|
| Control | F | 82 | 2.25 | 940 | II | |
| Control | M | 91 | 1.5 | 1440 | III | |
| Control | M | 82 | 2.16 | 1160 | II | 0 |
| Control | M | 78 | 2.75 | 1200 | II | 0 |
| Control | M | 97 | 1.5 | 1260 | III | 0 |
| ***Mean*** | ***4/1*** | ***86*** | ***2.03*** | ***1200*** | **-** | **-** |
| PD | F | 82 | 2.75 | 1215 | IV | III |
| PD | F | 73 | 2.16 | 1170 | IV | IV |
| PD | M | 85 | 2.16 | 1320 | III | IIb |
| PD | F | 84 | 2.0 | 1220 | II | IIa |
| PD | M | 72 | 3.5 | 1360 | II | III |
| ***Mean*** | ***2/3*** | ***79.2*** | ***2.51*** | ***1257*** | **-** | **-** |

**Table S2**. Demographic and clinical data of *post-mortem* putaminal tissue donors (Netherlands Brain Bank NBB)

| Diagnosis | Sex | Age (yrs) | Post mortem interval | Brain weight (g) | Braak NF score | Braak Lewy stage |
|---|---|---|---|---|---|---|
| Control | M | 77 | 507 | 1319 | II | 0 |
| Control | M | 77 | 435 | 1295 | I | 0 |
| Control | F | 81 | 444 | 1180 | II | 0 |
| Control | M | 92 | 435 | 1360 | III | 0 |
| Control | F | 80 | 507 | 1220 | III | 0 |
| ***Mean*** | ***3/2*** | ***81*** | ***466*** | ***1275*** | | |
| PD | M | 77 | 429 | 1285 | III | 6 |
| PD | F | 82 | 550 | 1162 | II | 6 |
| PD | M | 77 | 381 | 1355 | II | 6 |
| PD | F | 79 | 207 | 1027 | II | 5 |
| PD | M | 92 | 606 | 1400 | IV | 5 |
| ***Mean*** | ***3/2*** | ***81*** | ***434,6*** | ***1246*** | | |

**Table S3**. List of antibodies

| Target | Source | Company | Catalog Nr. | RRID | Application/ dilution |
|---|---|---|---|---|---|
| aSyn (Syn1) | Mouse | BD Biosciences | 610786 | AB_398108 | WB/ 1:1000 |
| aSyn (LB509) | Mouse | Abcam | ab27766 | AB_727020 | ICC/ 1:50 |
| aSyn (15G7) | Rat | Enzo Lifesciences | ALX-804-258-L001 | AB_2270759 | ICC/ 1:100 |
| aSyn (D1R1) | Rabbit | Cell Signaling | 23706 | AB_2798868 | WB/ 1:1000 |
| aSyn (MJFR-14-6-4-2) | Rabbit | Abcam | ab209538 | AB_2714215 | Immuno staining in filter trap assay/ 1:1000 |
| TUBB3 | Mouse | BioLegend | 801201 | AB_2313773 | ICC/ 1:500 |
| TUBB3 | Rabbit | BioLegend | 802001 | AB_2564645 | ICC/ 1:500 |
| TH | Rabbit | Millipore | AB152 | AB_390204 | ICC/ 1:300<br>WB/ 1:1000 |
| TH | Goat | Santa Cruz Biotechnology | Sc-7847 | AB_671396 | IHC/ 1:100 |
| FICD | Rabbit | Sigma-Aldrich | HPA021390 | AB_1851340 | ICC/ 1:200<br>IHC/ 1:400<br>WB/ 1:1000 |
| BiP | Rabbit | Thermo Fisher Scientific | PA5-34941 | AB_2552290 | WB/ 1:1000 |
| AMP-Thr | Mouse | Biointron | 17G6-1 | - | WB/ 1:1000 |
| CTSB | Goat | R&D Systems | AF953 | AB_355738 | WB/ 1:500 |
| GAPDH | Mouse | Millipore | MAB374 | AB_2107445 | WB/ 1:5000 |
| Mouse Alexa-647 | Donkey | Thermo Fisher Scientific | A-31571 | AB_162542 | WB/ 1:1000<br>ICC/ 1:500 |
| Mouse Alexa-568 | Donkey | Thermo Fisher Scientific | A10037 | AB_2534013 | WB/ 1:1000<br>ICC/ 1:500 |
| Mouse Alexa-488 | Donkey | Molecular Probes | A-21202 | AB_141607 | WB/ 1:1000<br>ICC/ 1:500 |
| Rabbit Alexa-647 | Donkey | Jackson ImmunoResearch | 711-605-152 | AB_2492288 | WB/ 1:1000<br>ICC/ 1:500 |
| Rabbit Alexa-568 | Donkey | Thermo Fisher Scientific | A10042 | AB_2534017 | WB/ 1:1000<br>ICC/ 1:500 |
| Rabbit Alexa-488 | Donkey | Molecular Probes | A-21206 | AB_2535792 | WB/ 1:1000<br>ICC/ 1:500 |
| Rat Alexa-568 | Goat | Thermo Fisher Scientific | A-11077 | AB_2534121 | WB/ 1:1000<br>ICC/ 1:500 |
| Rabbit IRDye 800CW | Donkey | Licor | 926-32213 | AB_621848 | WB/ 1:10000 |
| Mouse IRDye 680RD | Donkey | Licor | 926-68072 | AB_10953628 | WB/ 1:10000 |

| | | | | | |
|---|---|---|---|---|---|
| Goat IRDye 800CW | Donkey | Licor | 926-32214 | AB_10153047 | WB/ 1:10000 |
| Goat-Cy3 | Donkey | Jackson ImmunoResearch Labs | 705-165-147 | AB_2340412 | IHC/ 1:800 |
| Rabbit-Cy5 | Donkey | Jackson ImmunoResearch Labs | 711-175-152 | AB_2617154 | IHC/ 1:800<br>IHC/ 1:1000 |

**Table S4. List of primers**

| Target | Primer pair | Reference |
|---|---|---|
| *FICD* | F: TGTGCTCAAAGGCCTCTACC<br>R: GCTTCCAACTTACCCGCTG | NM_007076.3 |
| *SNCA* | F: CCATGGATGTATTCATGAAAGGACT<br>R: AAGTGGTCGTTGAGGGCAATG | NM_000345.4 |
| *18S* | F: GGAGTATGGTTGCCAAGCTGA<br>R: ATCTGTCAATCCTGTCCGTGT | (3) |
| *GAPDH* | F: CTGGGCTACACTGAGCACC<br>R: AAGTGGTCGTTGAGGGCAATG | NM_002046.7 |
| *ATF6* | F: CAGACAGTACCAACGCTTATGCC<br>R: GCAGAACTCCAGGTGCTTGAAG | NM_001410890.1 |
| *BiP* | F: GGAAAGAAGGTTACCCATGCA<br>R: GAGACACATCGAAGGTTCCG | NM_005347.5 |
| *PERK (EIF2AK3)* | F: GTCCCAAGGCTTTGGAATCTGTC<br>R: CCTACCAAGACAGGAGTTCTGG | NM_001313915.2 |
| *IRE1a (ERN1)* | F: CCGAACGTGATCCGCTACTTCT<br>R: CGCAAAGTCCTTCTGCTCCACA | NM_001433.5 |
| *CHOP (DDIT3)* | F: AATGAACGGCTCAAGCAGGA<br>R: AGCCACTTCTGGGAAAGGTG | NM_001195053.1 |
| *uXBP1* | F: CAGACTACGTGCACCTCTGC<br>R: CTGGGTCCAAGTTGTCCAGAAT | (4) |
| *sXBP1* | F: TGCTGAGTCCGCAGCAGGTG<br>R: GCTGGCAGGCTCTGGGGAAG | (5) |

### Supplementary methods

*Phos-tag analysis*

Phos-tag analysis of aSyn was performed as previously described (6), but after blotting aSyn was fixed with 4% PFA for 15 min at RT.

*Intact protein mass spectrometry of aSyn*

For top-down aSyn measurements, cell lysates were desalted on the ZipTip with C4 resin (Millipore, ZTC04S096) and eluted with 50 % (v/v) acetonitrile 0.1 % (v/v) formic acid (FA) buffer resulting in ~10 µM final protein concentration in 200–400 µl total volume. MS measurements were performed on an Orbitrap Eclipse Tribrid Mass Spectrometer (Thermo Fisher Scientific) via direct injection, a HESI-Spray source (Thermo Fisher Scientific) and FAIMS interface (Thermo Fisher Scientific) in a positive, peptide mode. Typically, the FAIMS compensation voltage (CV) was searched by a continuous scan. The most intense signal was usually obtained at -26 CV. The MS spectra were acquired with at least 120,000 FWHM, AGC target 100 and 2-5 microscans. The spectra were deconvoluted in Freestyle (Thermo) using the Xtract Deconvolution algorithm.

*Lentivirus production*

Third-generation lentiviral particles were produced following a previously established protocol (7, 8). The expression vector was modified by cloning the coding sequences for FICD variants (WT, H363A, or E234G) upstream of the internal ribosome entry site (IRES) and the GFP reporter. HEK293T cells were transfected with the expression construct and packaging plasmids to induce viral particle assembly. Viral supernatants were harvested 48 h post-transfection and clarified by filtration through a 0.45 µm PVDF membrane. Finally, the viral particles were pelleted by ultracentrifugation at 28,000 rpm for 2 h at 4°C. After centrifugation, the pellet was resuspended in ice-cold DMEM over night at 4°C and immediately stored at – 80°C.

*Lentiviral titer analysis via flow cytometry*

Functional lentiviral titer was determined by flow cytometry based on the fraction of GFP-positive HEK293T cells after transduction with serial viral dilutions. On day 1, 100.000 cells were seeded per well in 6-well plates in IMDM medium (Thermo Fisher Scientific) supplemented with 10% FCS. On day 2, viral supernatant was serially diluted in IMDM + 10% FCS to generate $10^{-1}$ to $10^{-5}$ dilutions and a medium-only control was included. Culture medium was aspirated and cells were incubated with 1 mL of the respective dilution per well for 48 h. On day 4, cells were harvested with 300 µL trypsin/EDTA for 3 min at RT, and then transferred into flow cytometry tubes. Cells were pelleted at 300 g for 5 min, fixed in 4% PFA for 15 min at RT, washed twice with PBS, resuspended in 300 µL PBS, and analyzed by flow cytometry (≥10,000 events per sample).
